# Dating *Alphaproteobacteria* evolution with eukaryotic fossils

**DOI:** 10.1101/2020.09.08.285460

**Authors:** Sishuo Wang, Haiwei Luo

## Abstract

Elucidating the timescale of evolution of bacteria is key to testing hypotheses on their co-evolution with eukaryotic hosts, which, however, is largely limited by the scarcity of bacterial fossils. Here, we incorporate eukaryotic fossils to date the divergence times of *Alphaproteobacteria*, based on the endosymbiosis theory that mitochondria evolved from an alphaproteobacterial lineage. We estimate that *Alphaproteobacteria* arose ~1900 million years (Ma) ago, followed by rapid divergence of their major clades. We show that the origin of *Rickettsiales*, an order of obligate intracellular bacteria whose hosts are mostly animals, predates the emergence of animals for ~700 Ma but coincides with that of eukaryotes. This, together with reconstruction of ancestral hosts, strongly suggests that early *Rickettsiales* lineages had established previously underappreciated interactions with unicellular eukaryotes. Our mitochondria-based approach displays higher precision and robustness to uncertainties compared with the traditional strategy using cyanobacterial fossils, and suggests that previous applications using divergence times of the modern hosts of symbiotic bacteria to date the bacterial tree of life may need to be revisited.

---

The *Alphaproteobacteria* is one of the largest grsoups within bacteria (Ettema and Andersson, 2009) and of great evolutionary significance for holding the origin of the mitochondrion (Wang and Wu, 2015; Yang et al., 1985). The *Alphaproteobacteria* has extensively diversified since its ancient origin, and comprised some of the most environmentally abundant and metabolically diverse organisms on Earth (Boussau et al., 2004; Ettema and Andersson, 2009; Luo et al., 2013). The intimate association between some alphaproteobacterial lineages and eukaryotes is of central importance for agricultural (e.g., rhizobia) and medical (e.g., rickettsia) applications. This makes *Alphaproteobacteria* a promising system to study the timing of bacterial evolution and their correlation with geological, ecological and evolutionary events (Luo et al., 2013; Sun et al., 2017; Wang et al., 2020). However, the traditional way of divergence time estimation has several limits when applied to *Alphaproteobacteria* and other prokaryotes. First, fossil records of bacteria are extremely scarce and controversial in terms of their estimated dates (Schirrmeister et al., 2016). Second, the most widely used prokaryotic fossils are from cyanobacteria, but the long evolutionary distance between cyanobacteria and other bacteria causes large uncertainties in dating (Nei et al., 2001). Last, some studies assumed a strict relationship of bacteria-host evolution, and calibrated the evolution of pathogenic/symbiotic bacteria based on the divergence time of their modern hosts (mostly animals and plants). However, this precludes the possibility of host switching, which could occur frequently during evolutionary processes spanning millions of years (Wang et al., 2020). Due to these challenges, the origin time of *Alphaproteobacteria* estimated by previous studies varies from less than 600 million years (Ma) to more than 2000 Ma (Battistuzzi et al., 2004; Battistuzzi and Hedges, 2009; Chriki-Adeeb and Chriki, 2016; Luo et al., 2013), making any narratives based on its evolutionary timing contentious.

Recently, horizontal gene transfer (HGT) has been suggested to have great potential in dating the evolution of bacteria (Davín et al., 2018; Wolfe and Fournier, 2018). Briefly, if the donor of an HGT event does not have fossil records while the recipient does, the temporal information recorded in the recipient can be transferred to date the evolution of the donor group (and vice versa), thereby bypassing the paucity of fossils in the donor lineage. Inspired by this idea, we developed a new strategy to date the divergence times of *Alphaproteobacteria* based on the endosymbiosis theory that the mitochondrion was transferred to a host nucleus from a bacterial lineage (Sagan, 1967), which was later shown to be within (Fan et al., 2020; Wang and Wu, 2015; Yang et al., 1985) or closely related to (Martijn et al., 2018) *Alphaproteobacteria* by modern phylogenetic analysis. As mitochondria are characteristic of eukaryotes, we took advantages of eukaryotic fossils to anchor the divergence time of *Alphaproteobacteria* in a tree integrating both alphaproteobacterial and mitochondrial lineages.

We first reconstructed a phylogenomic tree of 80 carefully selected *Alphaproteobacteria* and mitochondrial genomes using 24 conserved genes based on prior phylogenomics studies (Martijn et al., 2018; Muñoz-Gómez et al., 2019) (see Methods and Supplementary Note S1.1). We employed rigorous approaches to delineate phylogenetic artefacts caused by long branch attraction and compositional heterogeneity (see Methods), and obtained results consistent with recent studies where i) *Rickettsiales*, *Holosporales* and *Pelagibacterales* (SAR11) had independent origins (Muñoz-Gómez et al., 2019), and ii) mitochondria branched as a sister to *Alphaproteobacteria* (Martijn et al., 2018) (Fig. S1A). We also tested the impact of alternative topologies on dating (Fig. S1C; see below). We compiled two datasets to estimate the time divergences within the *Alphaproteobacteria* calibrated by eukaryotic fossils with relaxed molecular clock (Drummond et al., 2006), which accounts for substitution rate variations among branches. The first dataset, which we referred to as the mito-encoded dataset, was based on the aforementioned 24 conserved gene encoded by mitochondrial genomes (Dataset S1), and the mitochondrial lineages mainly comprised species of green plants, red algae and jakobids, whose mitochondrial genomes are both gene-rich and relatively slowly-evolving (Martijn et al., 2018; Rodríguez-Ezpeleta and Embley, 2012). Four high-confidence fossils from land plants and red algae were used as the calibration points (Supplementary Note S2.1; Fig. S2A). The second dataset, referred to as the nuclear-encoded dataset, was based on 22 mitochondria-derived genes that had been transferred to the nuclear genome identified by Wang and Wu, 2015 (Dataset S1; see Methods). This dataset not only circumvented the problem that in many eukaryotes genes encoded by mitochondrial genomes are few (e.g., apicomplexans and dinoflagellates) or fast-evolving (e.g., animals and fungi) (Roger et al., 2017), but integrated six additional eukaryotic fossils (Fig. S2A; Supplementary Note S2.1), allowing directly comparing the divergence time between symbiotic bacteria and their eukaryotic hosts on the same tree.

We selected a best-practiced scheme based on systematic comparisons of different combinations of parameters of MCMCTree for the mito- and nuclear-encoded datasets (see Methods and Supplementary Note S1.2). Similar divergence times were recovered for most nodes between the two datasets, although the mito-encoded dataset estimated older ages for deep nodes (Fig. 1). As shown in the infinite-sites plots (up- and bottom-left panels in Fig. 1), the posterior mean ages versus 95% HPD (highest posterior density) widths approached a straight line, suggesting that the uncertainty in time estimate was predominantly caused by the uncertainty associated with fossil calibrations (Rannala and Yang, 2007). The estimated ages in the nuclear-encoded dataset exhibited smaller 95% HPD intervals, hence smaller uncertainties, likely due to their more calibration information compared with the mito-encoded dataset.

**Figure 1.**
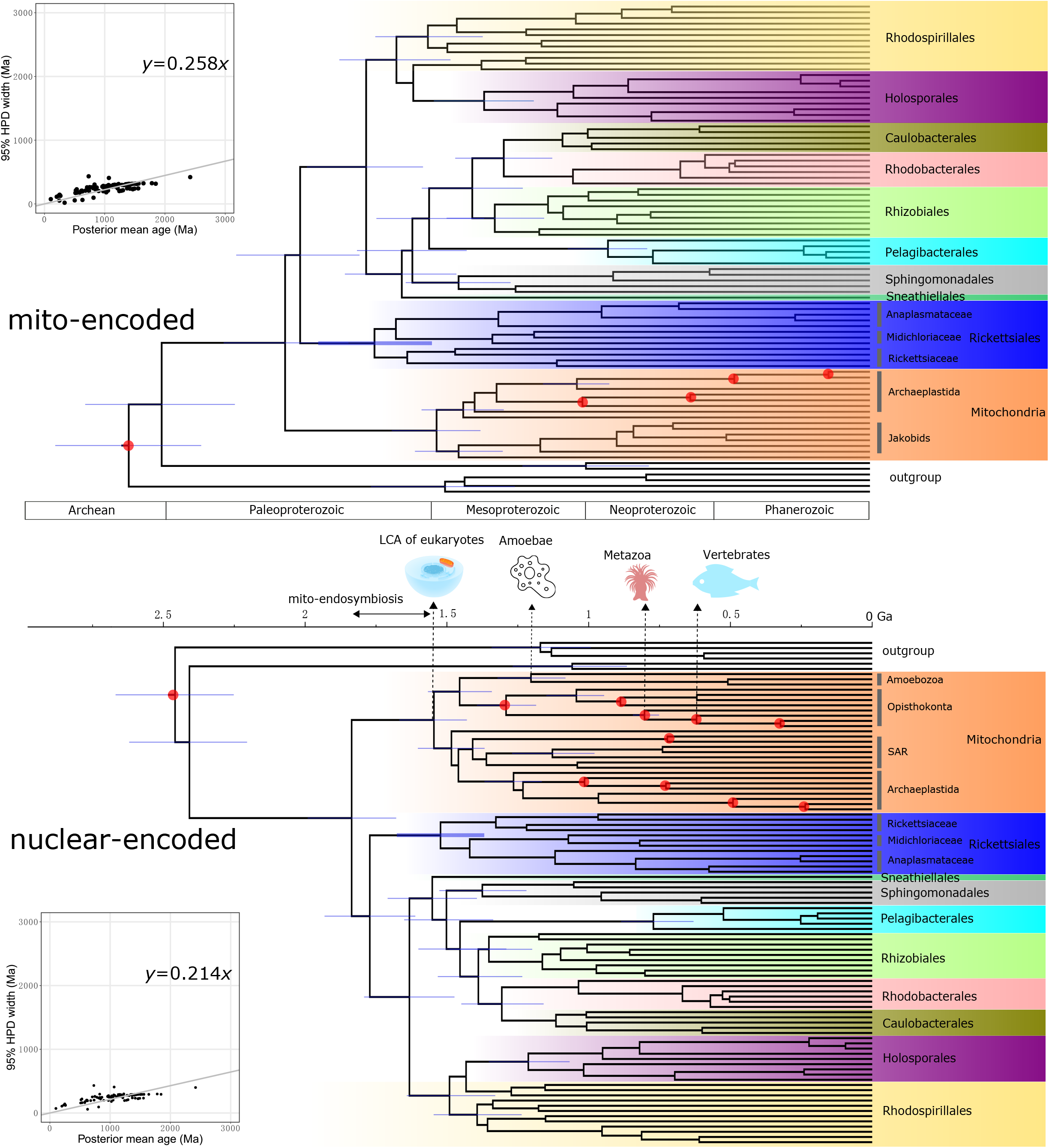
Divergence time estimate using the mitochondria-based strategy. The evolutionary timeline of the *Alphaproteobacteria* estimated using the best-practiced dating scheme for the mito-encoded (the top) and nuclear-encoded (the bottom) datasets. Node bars denote the 95% HPD interval of estimated dates. Nodes with red circles denote calibration points. The cartoon graphs above the timeline indicate the origin times of eukaryotic lineages that *Rickettsiales* and *Holosporales* are mostly associated with, inferred simultaneously with *Alphaproteobacteria* using the same dataset. The potential time range of mitochondrial endosymbiosis is also indicated. The panels in the upper and lower left corners are the infinite-sites plots, where the uncertainty in the divergence time (measured as the 95% HPD width) is plotted against the posterior mean of estimated times for each node. A lower value of the slope indicates less changes in the 95% HPD width, hence higher precision in dating. The sources and credits of the cartoon graphs are provided in the online open access repository FigShare (see Data availability).

Most alphaproteobacterial orders diverged 1500-1000 Mya, and *Rickettsiales* and *Pelagibacterales* appeared to be the oldest and youngest alphaproteobacterial orders, respectively (Fig. 1). The origin times of *Rickettsiales*, an obligate endosymbiont lineage whose hosts cover diverse eukaryotes but mostly animals (Merhej and Raoult, 2011), were estimated to be 1771 Ma (95% HPD 1972-1565 Ma) and 1523 Ma (95% HPD 1677-1368 Ma) using the mito- and nuclear-encoded datasets, respectively (Fig. 1). A merit of our mitochondria-based method is that divergence times of the eukaryotic hosts and of the host-associated bacteria can be simultaneously estimated. As shown in Fig. 1, we dated the origin of animals to be 803 Ma (95% HPD 844-752 Ma), consistent with previous dating analyses (Betts et al., 2018; Dos Reis et al., 2015; Parfrey et al., 2011), but not others (Hedges et al., 2004). We also estimated that mitochondrial lineages diverged from *Alphaproteobacteria* ~1900 Ma ago and that the last common ancestor (LCA) of mitochondria occurred ~1550 Ma ago (Fig. 1). Thus, the origin of *Rickettsiales* likely predated the evolutionary emergence of animals for ~700 Ma but coincided with the mitochondrial endosymbiosis process and the occurrence of the LCA of eukaryotes, according to our (Fig. 1) and others’ estimates (Betts et al., 2018; Eme et al., 2014; Parfrey et al., 2011), and fossil records (reviewed in Butterfield, 2015). This agrees with recent findings of an increasingly broad range of protistan hosts of *Rickettsiales* (reviewed in Castelli *et al.*, 2016), and suggests that host switches to animals from protists occurred later in the evolution of *Rickettsiales*. The origin time of *Holosporales*, another important endosymbiotic lineage in *Alphaproteobacteria* whose extant members are mostly endonuclear parasites of the ciliate *Paramecium* (Schrallhammer et al., 2018), dated back to 1379 Ma (95% HPD 1559-1201 Ma) or 1212 Ma (95% HPD 1353-1067 Ma), respectively, based on the mito-or nuclear-encoded datasets. This implies that the origin of *Holosporales* roughly coincided with the that of ciliates, which dated back to ~1150 Ma according to other estimates (Fernandes and Schrago, 2019; Parfrey et al., 2011). While the above analyses were based on amino acid sequences using MCMCTree, the basic patterns held similar with PhyloBayes or using coding sequences (Fig. S3).

We assessed the impact of uncertainties in Bayesian relaxed molecular clock time estimation, including the disparity between fossil evidence and molecular clock estimates, root age, across-branch rate variation, sequence partitioning, and clock model, on the posterior ages (Dataset S2). When we used only Phanerozoic fossils and excluded all Proterozoic fossils (which are thought controversial by some), the estimated ages of most nodes were shifted towards the present for the mito-encoded dataset but not for the nuclear-encoded dataset (*Phan* in Figs. 2A, S4). Removing potentially controversial maximum age constraints led to minor changes in the posterior ages (*Max-1* in Figs. 2A, S4). Using more conservative calibrations of the root pushed the time estimates to be slightly older for the mito-encoded dataset (*Root-1* in Figs. 2A, S4). Allowing larger rate variation among branches resulted in highly consistent results with the best-practiced scheme (*Sigma* in Fig. S5). Decreasing the number of partitions resulted in decreased precision, as indicated by the increase of the slope in infinite-sites plots (Fig. S6), but the estimated dates remained similar (*Single partition* in Fig. 2A). The largest changes in the posterior ages were obtained when the independent rates (IR) instead of autocorrelated rates (AR) clock model was used: the divergence times of most alphaproteobacterial orders were shifted towards the present by ~20% (*IR* in Figs. 2A, S4). Collectively, the composite of the ages estimated from six different analyses shows that *Alphaproteobacteria* originated 2012 Ma (95% HPD 2284-1750 Ma) and 1786 Ma (95% HPD 2004-1574 Ma), based on the mito- and nuclear-encoded datasets, respectively (Fig. S7), and diversified soon thereafter.

**Figure 2.**
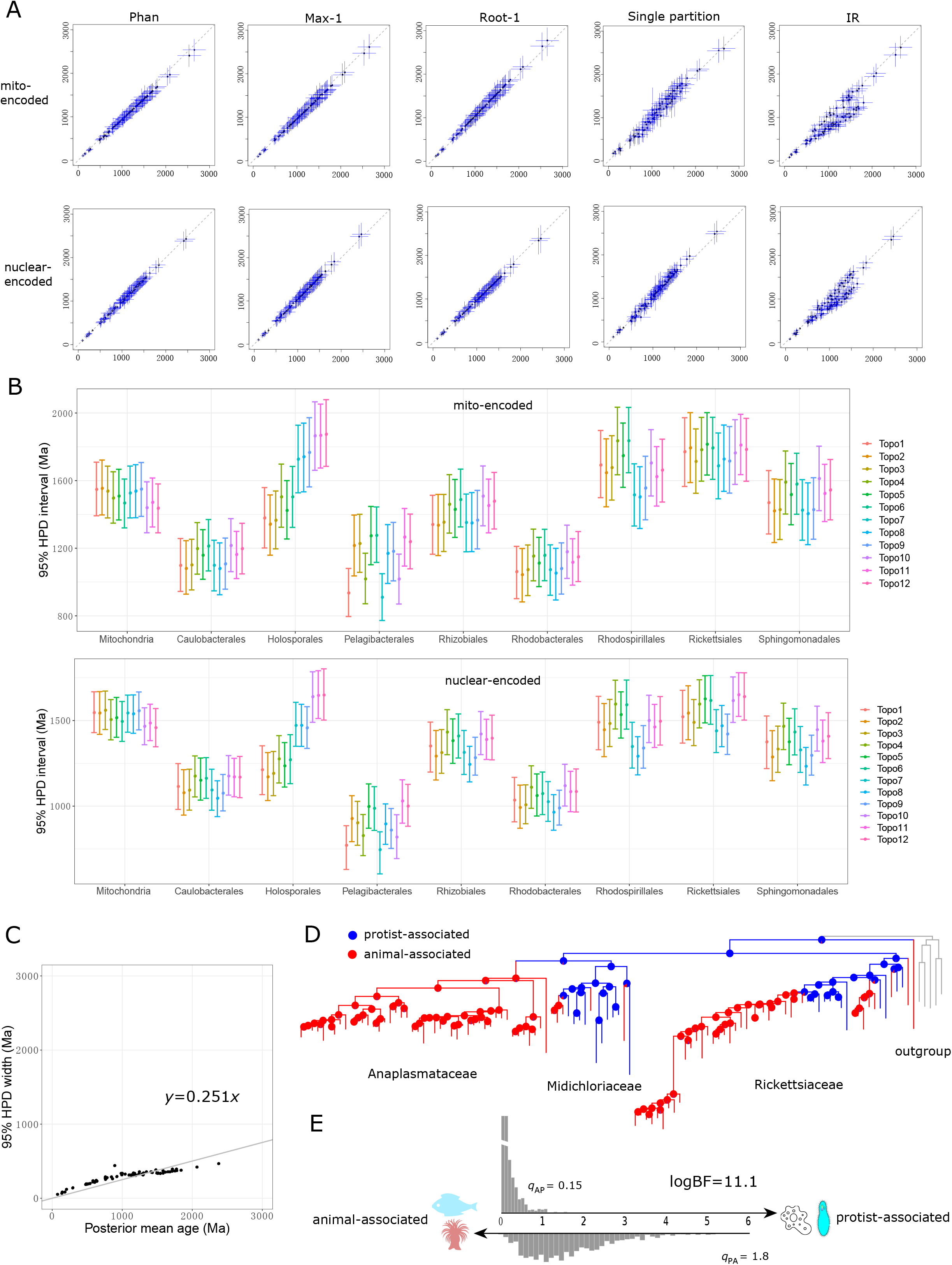
Comparison of the estimated times with alternative dating strategies using MCMCTree and ancestral lifestyle reconstruction of *Rickettsiales*. (A) The divergence times estimated using alternative schemes (y-axis; see Dataset S2) versus using the best-practiced scheme (x-axis). The best-practiced scheme used a full partition and autocorrelated rates clock model (Dataset S2). The bars in blue indicate the 95% HPDs. *Phan*: only calibration points with Phanerozoic fossils considered; *Max-1*: maximum constraints for nodes with controversial maximum ages removed; *Root-1*: a more conservative root age; *Single partition*: all sequences considered as a single partition; *IR*: independent rates clock model (Dataset S2). (B) Changes in the estimated times (95% HPD) that result from using different species tree topologies (Fig. S1C). The detailed posterior dates for each clade are shown in Table S2. (C) The infinite-sites plot using the cyanobacteria-based strategy. (D) Inferred ancestral hosts of *Rickettsiales*. The pie charts on the nodes show the estimated probabilities of the hosts, and the branch colors indicate the hosts with the higher probability at the corresponding node. Tips represent the randomly selected representative of each OTU (defined by 97% identity of 16S rRNA gene). (E) The transition rates from animal-associated to protist-associated (q_AP_) and from protist-associated to animal-associated (q_PA_) estimated by the MCMC method in BayesTraits multistate. The log-transformed Bayes factor (logBF) is indicated, where values above 10 are considered very strong evidence for support (Meade and Pagel, 2016). The sources and credits of the cartoon graphs are provided in the online open access repository FigShare (see Data availability).

The topology of *Alphaproteobacteria* is another much-debated issue (Fan et al., 2020; Luo, 2015; Muñoz-Gómez et al., 2019; Wang and Wu, 2015). We repeated the MCMCTree analysis by fixing the species phylogeny to 11 alternative topologies (Fig. S1B). Most alphaproteobacterial orders showed highly consistent estimated ages (Fig. 2B). However, the posterior mean ages of *Holosporales* varied from ~1800 to ~1200 Ma across different topologies. This was because the alternative phylogenetic position of *Holosporales* was a sister to *Rickettsiales*, in contrast to the topology used in the main analysis where *Holosporales* branched within the *Rhodospirillales* (Fig. S1B). Likewise, the origin time of *Pelagibacterales* showed considerable variations depending on whether it formed a monophyletic group with the *Rickettsiales*. On the other side, the impact of the topology of the eukaryotic tree (mitochondria subtree) was relatively minor, as different topologies showed similar estimates of alphaproteobacterial lineages at the order level (Fig. S8).

Traditionally, constructing the evolutionary timeline for bacteria is based on fossils of cyanobacteria since bacterial fossils that can be accurately assigned to a taxonomic group are only available for cyanobacteria (Schirrmeister et al., 2016). Based on a best-selected scheme, the dates estimated using calibrations within cyanobacteria were generally comparable to using the mitochondria-based strategy (Fig. S9). However, with a slope of 0.251 in the infinite-sites plot (Fig. 2C), the cyanobacteria-based strategy exhibited less precision compared with the nuclear-encoded dataset (slope of 0.214 in the infinite-sites plot in Fig. 1). Further, analyses with different calibration schemes within cyanobacteria revealed larger variations in the estimated ages for some clades (Fig. S9). For example, by using different calibration schemes, we observed a spread of 444 Ma for the posterior mean age of the *Rickettsiales* crown group with the cyanobacteria-based strategy, in contrast to 103 and 82 Ma as estimated by the mito-encoded and nuclear-encoded datasets of the mitochondria-based method, respectively (Fig. S9), likely underlined by the abundant high-confidence fossils in eukaryotes and the shorter phylogenetic distance between the mitochondrial lineage and *Alphaproteobacteria*.

The abundant eukaryotic fossils in our mitochondria-based strategy greatly reduced dating uncertainty compared with the cyanobacteria-based approach. By co-estimating the divergence times of both bacteria and their eukaryotic hosts, it allows directly analyzing their co-evolution with the same dataset on the same tree, thereby avoiding methodological artefacts stemming from comparing age estimates from different studies. In addition, the mitochondria-based approach to date evolution is conceptually distinct from the HGT-based method (Davín et al., 2018; Wolfe and Fournier, 2018), as mitochondrial endosymbiosis is far more complex than simply an HGT event (Roger et al., 2017; Sagan, 1967). Consequently, thanks to the many high-confidence orthologs shared by mitochondrial and alphaproteobacterial lineages, our taxon- and gene-rich multigene data mitigate the challenges faced by others (Shih et al., 2017) (see Supplementary Note S3.1). Nevertheless, our tentative analysis is subject to other challenges such as the lack of internal fossils within the *Alphaproteobacteria* and violation to molecular clock caused by the fast-evolving nature of mitochondrial genomes. Although its impact should be attenuated by the relaxed molecular clock model, it is important to keep this limit in mind when interpreting the results. Further, the mitochondria-based strategy works well for *Alphaproteobacteria*, but calibrating the evolution of related bacteria like *Beta-* and *Gammaproteobacteria* with this strategy requires additional testing and benchmarking.

The divergence times estimated here agree with some previous studies (Battistuzzi et al., 2004; Battistuzzi and Hedges, 2009), but are much older than in other studies that constrained the origin time of symbiotic or pathogenic bacteria to be the same as that of their dominant modern hosts, thus ignoring the possibility of host shifts (Chriki-Adeeb and Chriki, 2016; Luo et al., 2013; Weinert et al., 2009) (Supplementary Note S3.2). Our results suggest that the origin of *Rickettsiales*, where most extant members adapt to an animal-associated lifestyle, predated animals’ emergence for ~700 Ma but coincided with the origin of eukaryotes (Fig. 1). Moreover, ancestral lifestyle reconstruction using BayesTraits (Meade and Pagel, 2016) with a much broader taxon sampling of *Rickettsiales* suggests the LCA of *Rickettsiales* to be symbionts of protists and that the transition rate from protist-associated to animal-associated lifestyle was more than ten times higher than that of the reverse process (Fig. 2D-E). This strongly challenges the view of the concurrence of *Rickettsiales* and animals (Weinert et al., 2009), and instead suggests frequent host transitions from unicellular eukaryotes to animals during *Rickettsiales* evolution. Presumably, the predatory nature of early eukaryotes like amoebae or ciliates might have helped forming primitive association with ancestral *Rickettsiales* (Vannini et al., 2005). Later, animals could have acquired *Rickettsiales* parasites early in their evolution by filter-feeding on infected protists (Supplementary Note S3.3). This scenario is also supported by recent findings of the association between *Rickettsiales* members with diverse protists (Castelli et al., 2016; Montagna et al., 2013) and by the presence of amoebae-derived genes in some *Rickettsiales* genomes (Ogata et al., 2006). *Holosporales* are mostly intracellular parasites of the ciliate *Paramecium*, but are increasingly found associated with a diverse range of protists including amoebae, rhizarians and *Euglenozoa* (Muñoz-Gómez et al., 2019), consistent with their deep divergence. Overall, our results are not contradictory to the idea of host-bacteria co-evolution, but suggest more frequent host transitions than previously understood and that caution should be exercised when assigning the divergence time of bacteria based on that of their modern hosts. Besides, the ubiquity of microbial eukaryotes like amoebae in modern environments reminds people to pay more attention to them as potential reservoirs of emerging human diseases.

## Methods

### Phylogenomic reconstruction

We followed Martijn *et al.*, 2018, and used the 24 conserved genes encoded by both the mitochondrial and alphaproteobacterial genomes annotated by MitoCOGs (Kannan et al., 2014) to determine their phylogenetic relationship. Eighty genomes were carefully selected for phylogenomic reconstruction based on prior studies (Supplementary Note S1.1; Table S1). Genes were aligned using MAFFT v7.222 (Katoh and Standley, 2013) and trimmed with TrimAl v1.4 (“-st 0.001”) (Capella-Gutiérrez et al., 2009). Because *Alphaproteobacteria* is subjected to strong compositional heterogeneity across lineages (Luo, 2015; Martijn et al., 2018; Muñoz-Gómez et al., 2019), which might confound phylogenetic signals and cause phylogenetically unrelated species with similar GC content to cluster together, we recoded the 20 amino acids into four nucleic acid characters according to their physicochemical properties with the dayhoff4 and SR4 recoding scheme, respectively (Luo, 2015; Martijn et al., 2018; Muñoz-Gómez et al., 2019). Phylogenomic reconstruction was performed under the empirical profile mixture model GTR+G+F+C30 with the PMSF approximation (guide tree GTR+G+I+F) and 1,000 ultrafast bootstraps using IQ-Tree v1.6.11 (Nguyen et al., 2015). As the trees obtained by dayhoff4 and SR4 recoding schemes showed similar topologies (Fig. S1), we used the one obtained by the dayhoff4 recoding scheme for dating (and we subsequently included more eukaryotic taxa as mitochondrial lineages based on the general consensus understanding of the eukaryotic phylogeny for both the mito- and nuclear-encoded datasets in dating [see Fig. S2A and Supplementary Note S1.1]).

### Calibration information

Four and ten calibration points within the eukaryote clade were selected for the mito- and nuclear-encoded datasets, respectively (Fig. S2). We based the lower limit of a calibration point upon the most ancient uncontroversial fossil from within the clade. Since fossil records only tell the time that the group of interest had already appeared, the actual origin time of a clade could be more ancient than these minima. Maximum time constraints were determined from the youngest geological formation or stratigraphic range without any members of the clade of interest, as used and recommended by many studies (Betts et al., 2018; Donoghue and Benton, 2007; Dos Reis et al., 2015). Several alternative time constraints were also considered to accommodate the uncertainties in calibration. The full details of calibrations are given in Supplementary Note S2.

### Divergence time estimation

We compiled two datasets for the mitochondria-based divergence time estimation of *Alphaproteobacteria*, the mito-encoded dataset and the nuclear-encoded dataset. The mito-encoded set was based on the aforementioned 24 orthologs conserved in *Alphaproteobacteria* and mitochondrial lineages. For the nuclear-encoded set, we retrieved the 29 genes that are likely transferred from the mitochondrial genome to the nuclear genome during the early evolution of eukaryotes from Wang and Wu, 2015, and excluded seven genes involving putative paralogs (Dataset S1). Wrongly annotated sequences in each alignment were removed by manually checking the alignment and gene phylogeny.

To alleviate the impacts of mutational saturation, we used amino acids in the main analysis but also repeated the analysis with nucleotide sequences (only the first two codon positions). Dating analyses were predominantly carried out with the approximate likelihood calculation with MCMCTree 4.9j (Yang, 2007), and we also examined the consistency of the results with PhyloBayes v4.1b (Lartillot et al., 2009). A constraint tree constructed using the dayhoff4 recoding scheme described above was applied as both MCMCTree and PhyloBayes require a fixed phylogeny topology. Because previous studies often came to different topologies of the *Alphaproteobacteria* phylogeny, which are mainly associated with the positions of mitochondria, *Holosporales* and *Pelagibacterales* (Fan et al., 2020; Luo, 2015; Martijn et al., 2018; Muñoz-Gómez et al., 2019), we considered two, two, and three distinct topologies for these three orders respectively, totaling 2 × 2 × 3 = 12 topologies (Fig. S1B; Supplementary Note S1.2). We addressed the topology uncertainty by repeating MCMCTree analysis with each alternative topology. We further selected a best-practiced dating scheme by investigating the impact of the calibration information, clock model, number of partitions, and cross-lineage rate variation on the estimated posterior ages (see Supplementary Note S1.2 for details).

The burn-in, sampling frequency, and number of the iterations were adjusted to 100,000, 100, and 20,000, respectively, based on the results of testing runs. This ensured that the effective sample size for all parameters were above 200, as commonly recommended for MCMC-based Bayesian phylogenetic inference (Nascimento et al., 2017). Convergence was assessed by comparing the posterior means from two independent chains and with Tracer v1.6 (http://tree.bio.ed.ac.uk/software/tracer/). The posterior ages were compared with effective priors (“usedata = 0”) to ensure that their distributions were different and thereby the sequences used in MCMCTree analysis were informative (Table S2). Further, we followed the above procedure to date the divergence time of *Alphaproteobacteria* using the traditional strategy where all calibration points were placed within cyanobacteria for comparison (see Supplementary Note S2.2).

## Data availability

All of the sequences, phylogenetic trees, molecular dating analysis results, and the Ruby (Goto et al., 2010) codes generating them, are available at https://figshare.com/s/ad430d1f5bace9eb2000.

## Acknowledgements

We are particularly grateful to Jan Janouškovec from University of Oslo for his insights on endosymbiosis and constructive comments on the draft of the manuscript. We thank Mario dos Reis from QMUL, Richard Brown from LJMU, Charles Foster from USYD, and Joana Wolfe from MIT for guidance in molecular dating, and Sergio Muñoz-Gómez from Dalhousie University for help in phylogenomics analysis. We thank lab members Tianhua Liao and Hao Zhang for discussion, and Kwok Chu Cheung for data retrieving.

This work is supported by the National Key R&D Program of China (2018YFC0309800), the Hong Kong Research Grants Council Area of Excellence Scheme (AoE/M-403/16), the Direct Grant of CUHK (4053257 & 3132809) and The CUHK Impact Scheme Fellowship to (S.W.).

