## Supplementary material for "Dating *Alphaproteobacteria* evolution with eukaryotic fossils": Figures S1-S12

Figure S1. Phylogenomic reconstruction of *Alphaproteobacteria* and mitochondria, and presentation of alternative topologies which differ in the phylogenetic positions of *Rickettsiales*, *Holosporales*, *Pelagibacterales*, and mitochondria. (A, B) The trees were generated using IQ-Tree under the C30 profile mixture model with PMSF approximation based on the dayhoff4 (A) and SR4 (B) recoding scheme, respectively. Circles in black and grey on the nodes denote nodes with the IQ-Tree ultrafast bootstrap values ≥ 95% and ≥ 85%, respectively. The scale bars denote 0.05 substitutions per amino acid site. The phylogeny shown in panel A was used in the main analysis for dating (with more eukaryotes added to the mitochondria subtree [see Fig. S2A and Supplementary Note S1.1]). (C) The 12 different topologies of the *Alphaproteobacteria* phylogeny used in dating. *Topo1* represents the one shown in panel A.

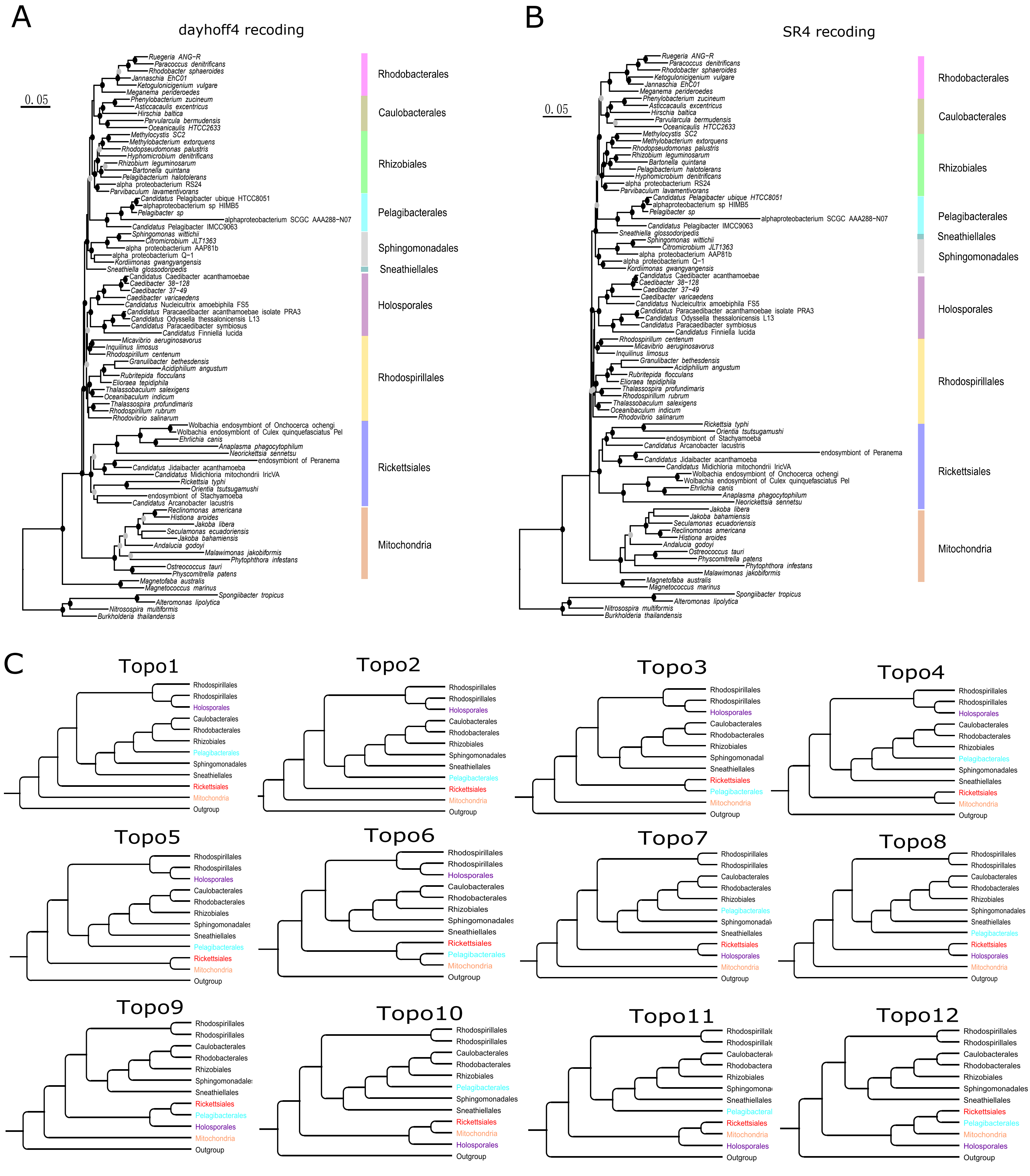

Figure S2. Calibration points for the mitochondria-based (A) and cyanobacteria-based (B) strategy. (A) Calibration information of all of the ten nodes were used in the dating analysis of the nuclear-encoded dataset. Only Nodes 1-4 were used for the mito-encoded dataset. The calibrations are detailed in Supplementary Note S2.1. Tips labelled with dark green and purple circles indicate those used in both datasets and only the nuclear-encoded dataset, respectively. Tips with a light green circle denote those used only in the mito-encoded dataset due to the unavailability of their nuclear genome sequences. The topology is based on general consensus understanding of the eukaryotic phylogeny (see Supplementary Note S1.1). (B) All of the three calibration points were involved in the best-practiced dating scheme (but not necessarily in alternative dating schemes) for cyanobacteria-based dating. The tree topology is according to IQ-Tree phylogenomic reconstruction. All species present in the tree were used in the cyanobacteria-based dating.

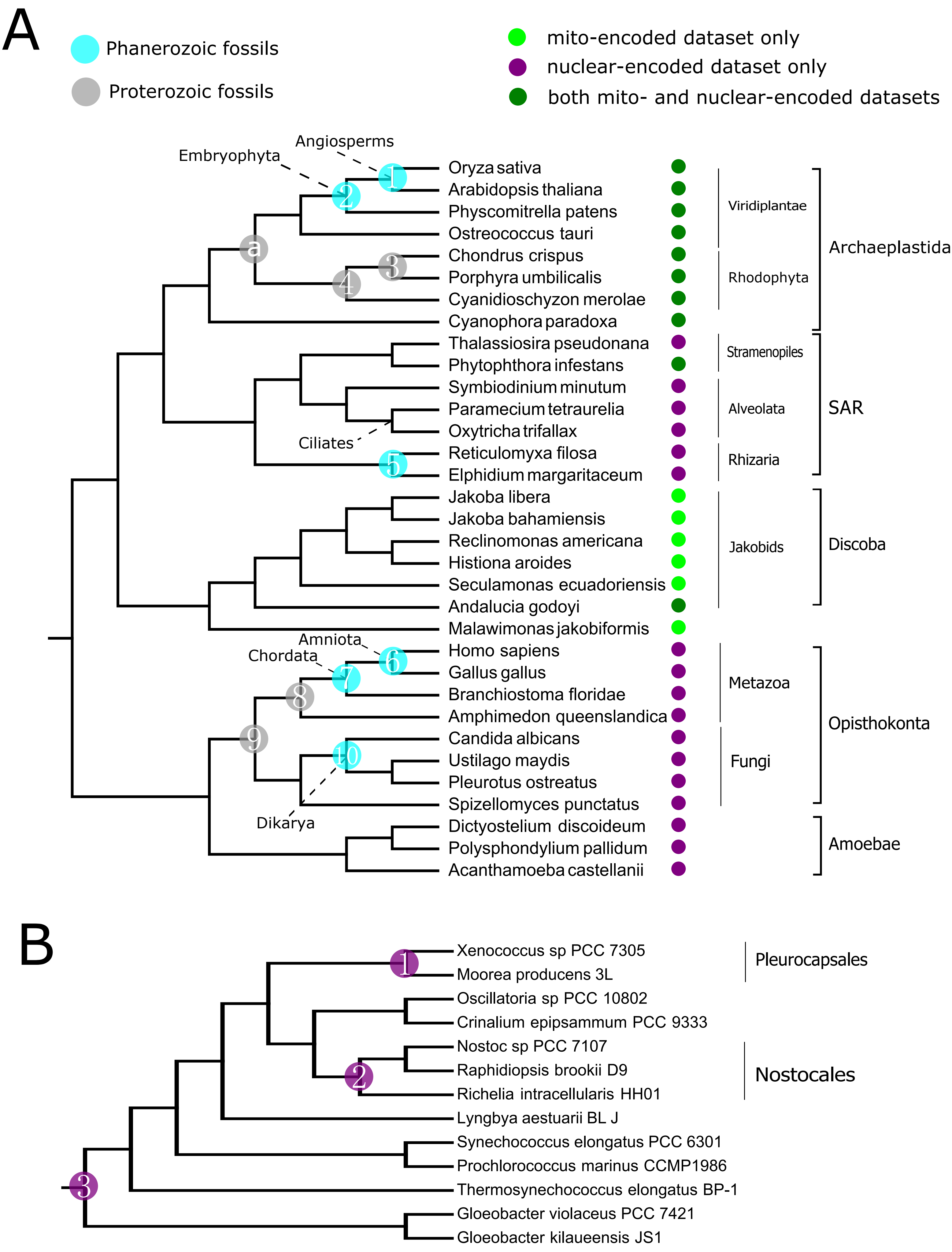

Figure S3. Comparison of the chronograms obtained using PhyloBayes with amino acid sequences or MCMCTree with nucleotide sequences and the best-practiced dating scheme for mito- and nuclear-encoded datasets. Nodes are drawn at the posterior means. (A) PhyloBayes with amino acid sequences and the log-normal autocorrelated clock model ("-ln"). (B) PhyloBayes with amino acid sequences and the uncorrelated gamma multipliers ("-ugam"). (C) MCMCTree with coding sequences (only the first two positions of a codon). (D) MCMCTree with amino acid sequences based on the best-practiced scheme.

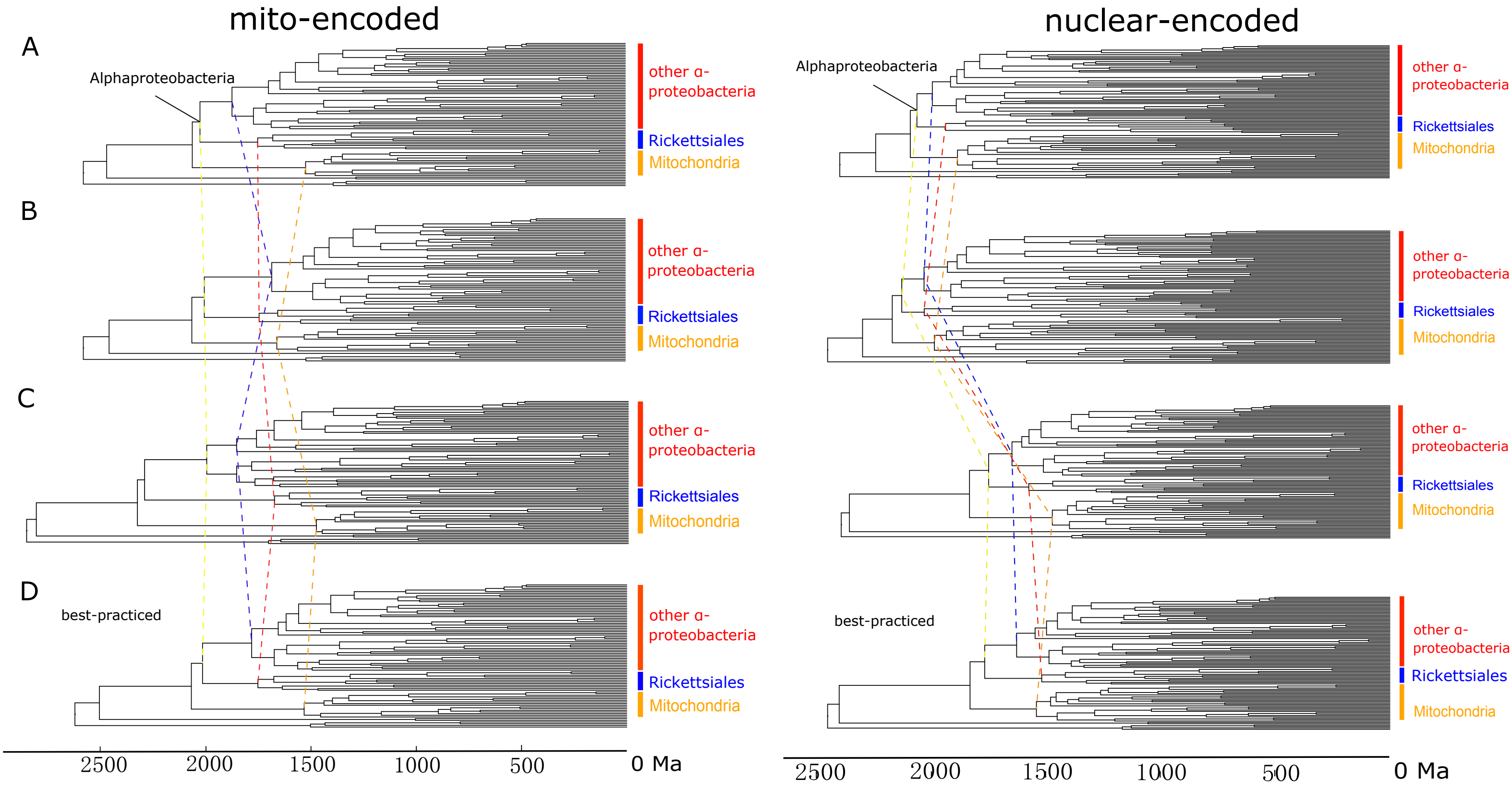

Figure S4. Comparison of the chronograms obtained with different dating schemes using MCMCTree for mito- and nuclear-encoded datasets. *Phan*: only calibration points with Phanerozoic fossils considered. *Max-1*: maximum constraints for nodes with controversial maximum ages removed. *Root-1*: a more conservative root age. *Single partition*: all sequences considered as a single partition. *IR*: independent rates clock model.

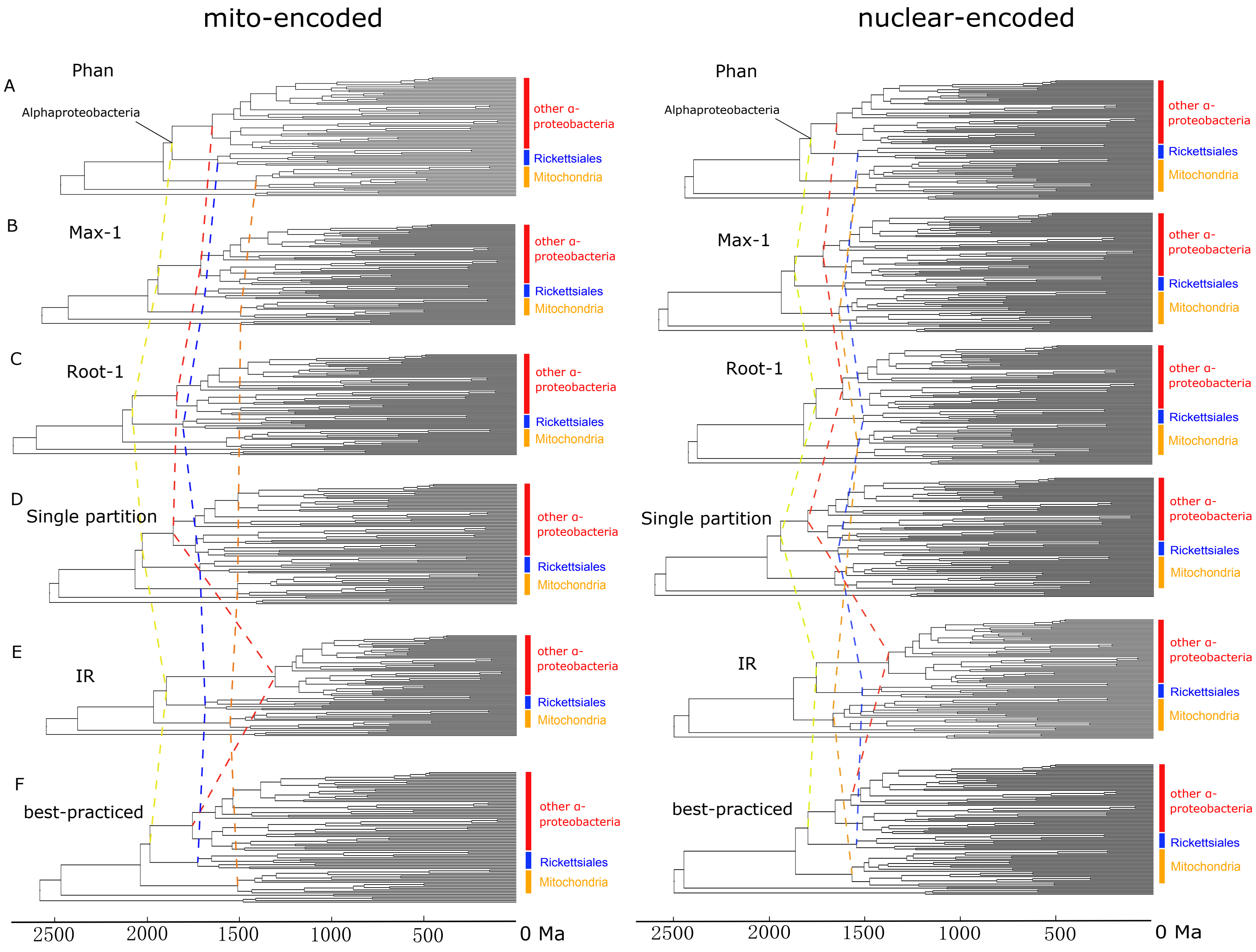

Figure S5. The divergence times estimated using alternative schemes (y-axis) versus using the best-practiced scheme (x-axis) using MCMCTree for the mito- and nuclear-encoded datasets (see Dataset S2 for details). *Max-2*: calibration points supported by only the minimum constraint were assigned with a maximum constraint of 1891 Ma (see Dataset S2), the earliest record of eukaryotes as used in the studies (Betts et al., 2018; Morris et al., 2018). *Root-2*: Root age calibrated as 3000-1500 Ma (uniform distribution). *Root-3*: Root age calibrated as 3000-2000 Ma (uniform distribution). *Sigma*: the prior of the standard deviation of log rate on branches was set to 1.0 to inform very large rate variation among branches.

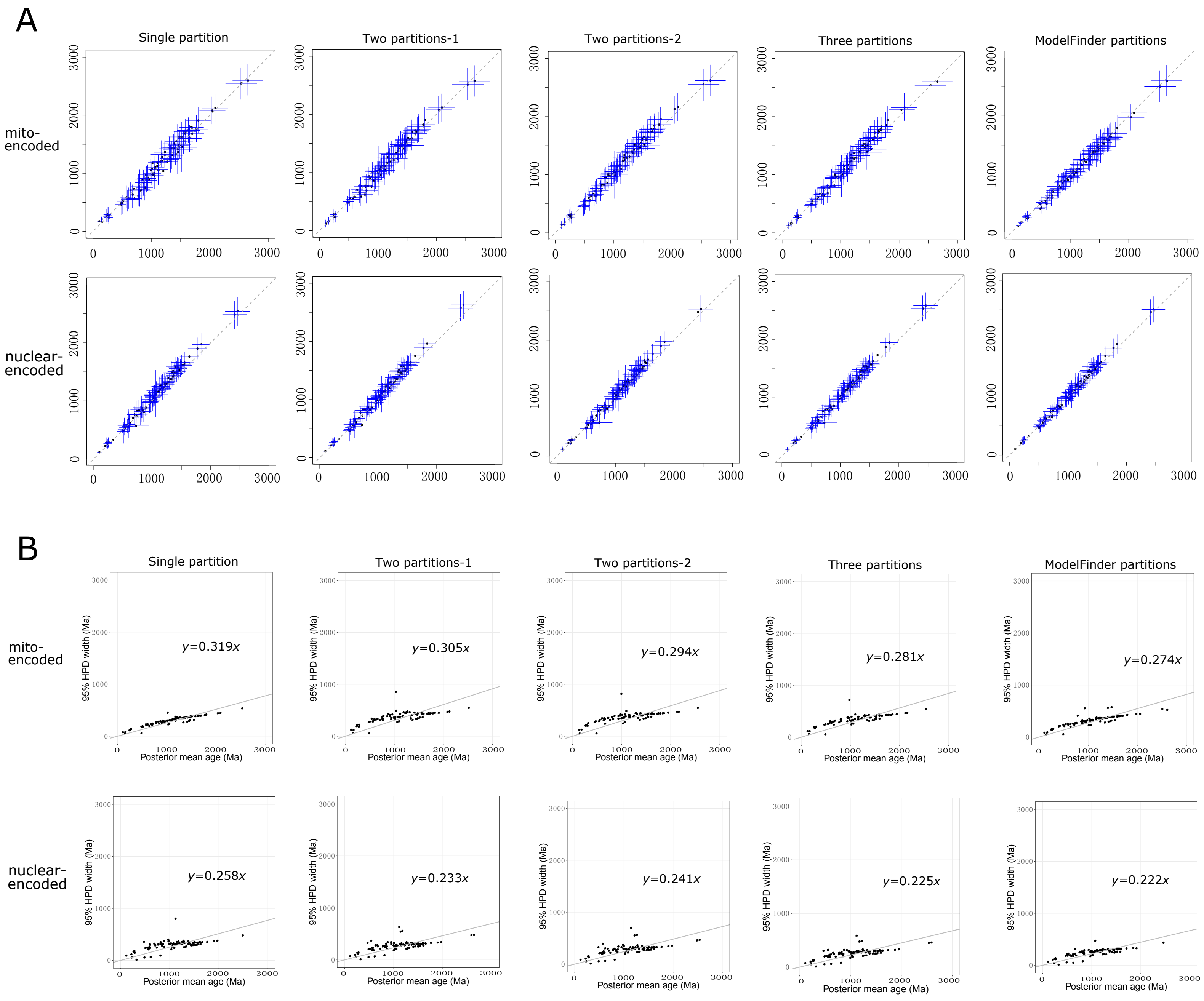

Figure S6. The divergence times (A) and infinite-sites plots (B) estimated with different sequence partitioning strategies in MCMCTree analysis for the mito-encoded and nuclear-encoded datasets. (A) The divergence times estimated using different numbers of partitions (y-axis) versus using the best-practiced scheme (fully partitioned) with MCMCTree. ModelFinder partitions indicate partitioning based on ModelFinder. Details of the partitioning are given in Dataset S2 and the online repository Figshare (see Data availability). (B) Infinite-sites plots obtained by using different numbers of partitions. The uncertainty in the divergence time, measured as the 95% HPD width, is plotted against the posterior mean of times for each node. A lower value of the slope indicates less changes in the 95% HPD width, hence higher precision in dating. The infinite-sites plots for the best-practiced dating scheme are displayed in Fig. 1.

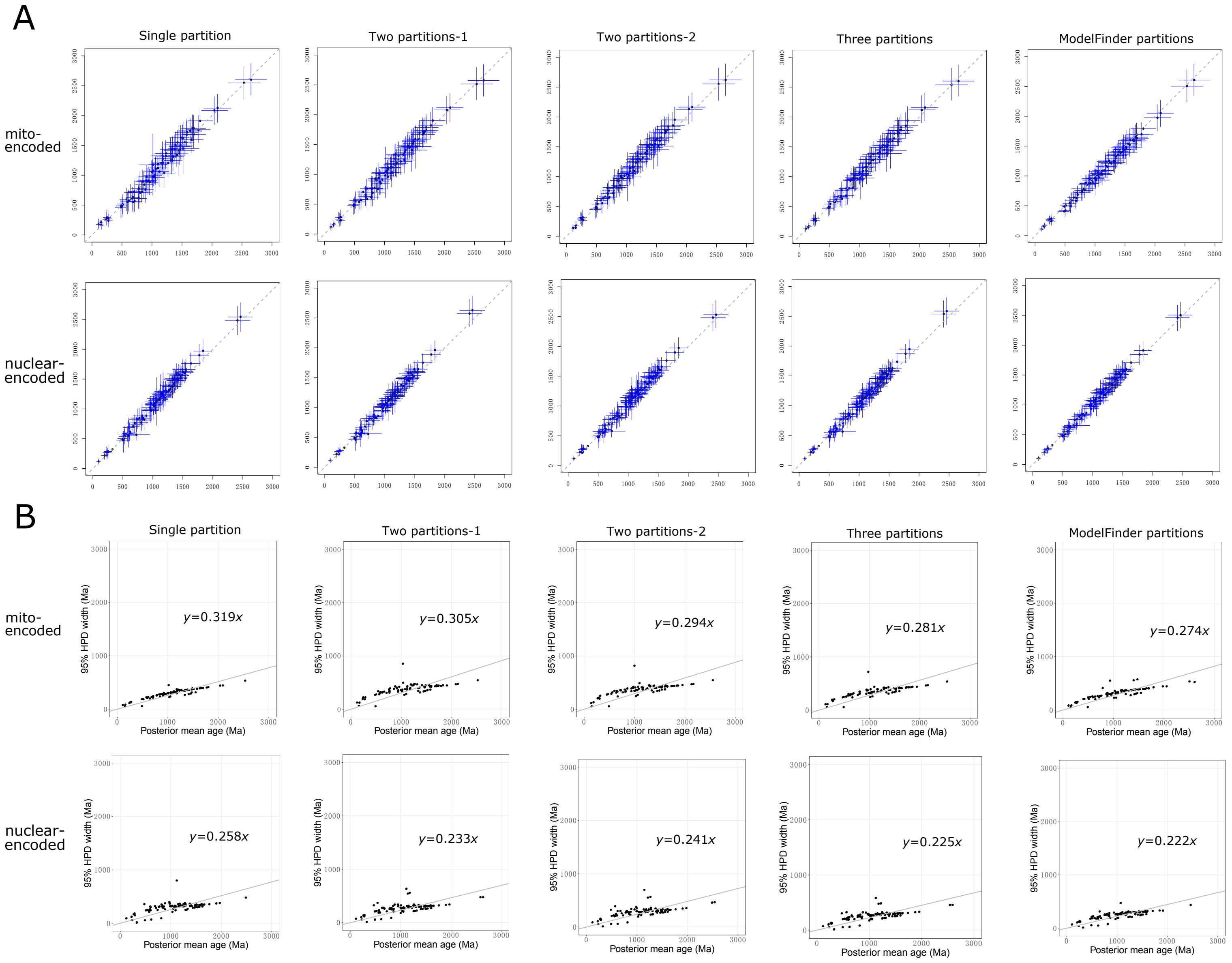

Figure S7. The composite chronograms combining uncertainties associated with Bayesian relaxed molecular clock analysis including calibrations, models, and partitions for the mitochondria-based strategy. The node ages and the 95% HPD intervals were derived from the composite of the posterior estimates across six competing dating schemes (the best-practiced scheme together with the five alternatives shown in Fig. 2A, namely *Phan*, *Max-1*, *Root-1*, *Single partition*, and *IR*). Node bars indicate 95% HPD intervals of posterior ages. (A) The mito-encoded dataset. (B) The nuclear-encoded dataset.

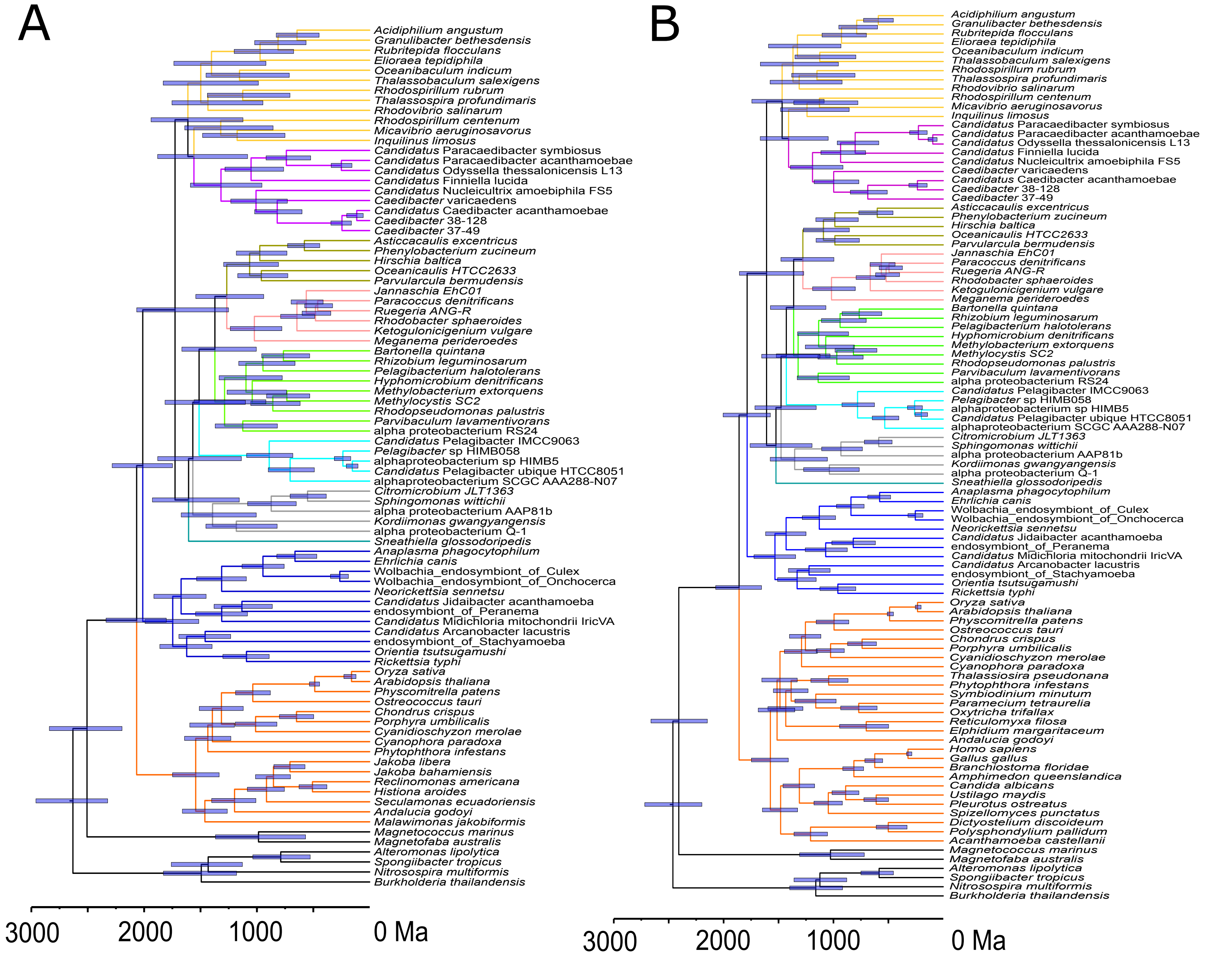

Figure S8. Comparison of the chronograms obtained with different topologies of the mitochondria subtree (shown in the left panel) using MCMCTree for the nuclear-encoded dataset.

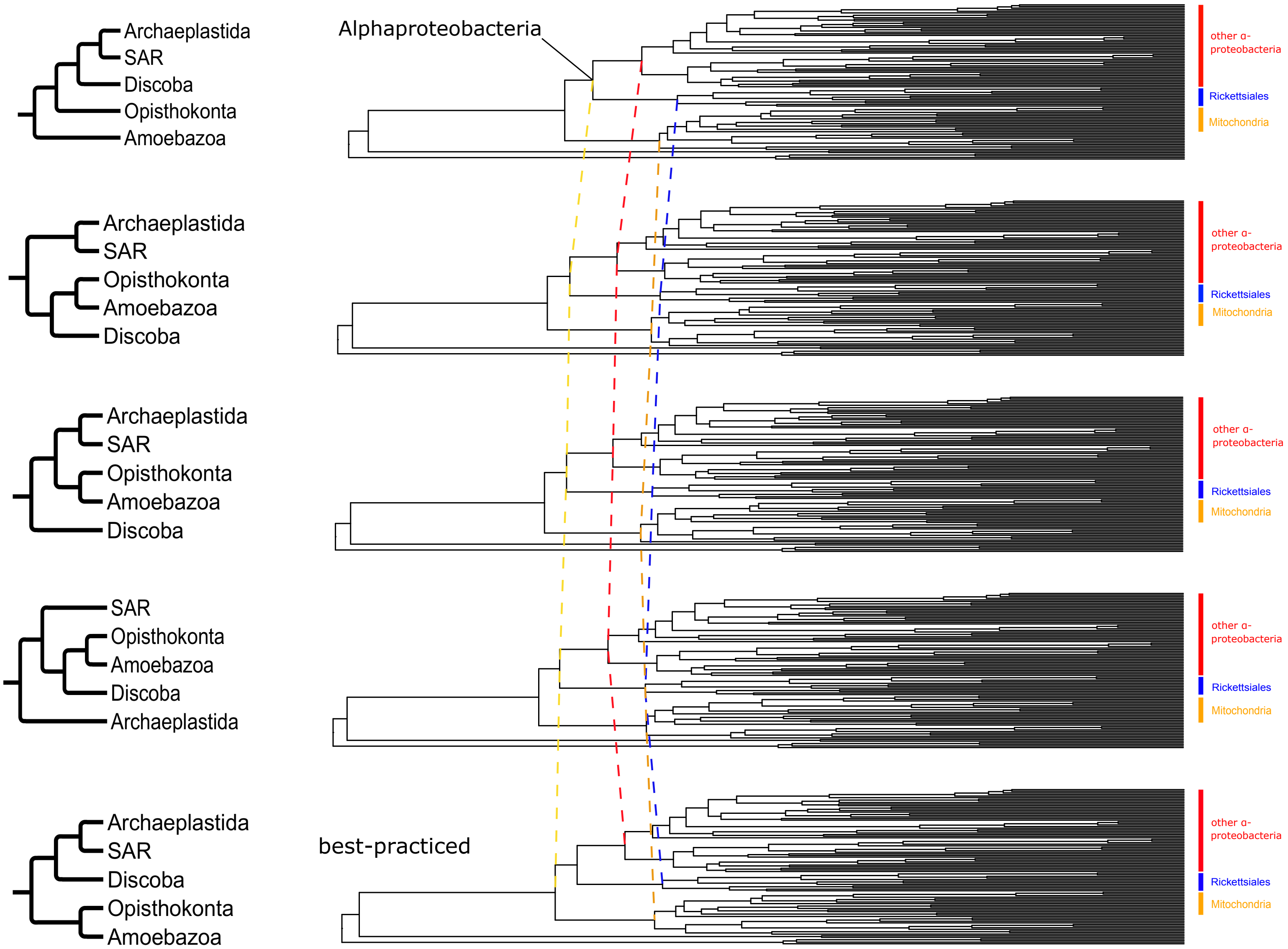

Figure S9. Comparison of the posterior ages of major alphaproteobacterial orders between the mitochondria- and cyanobacteria-based strategies. The bars indicate the 95% HPD intervals. The dots on the lines indicate the means. Different colors of the bars represent different calibration schemes (see Dataset S2). For the mito- and nuclear-encoded datasets: *Phan*: only calibration points with Phanerozoic fossils considered. *Max-1*: maximum constraints for nodes with controversial maximum ages removed. *Max-2*: calibration points supported by only the minimum constraint were assigned with a maximum constraint of 1891 Ma (see Dataset S2), the earliest record of eukaryotes as used in the studies (Betts et al., 2018; Morris et al., 2018). *Root-1*: a more conservative root age. For the cyanobacteria-based strategy: *Sanchez 2014*: calibrations according to Sánchez-Baracaldo *et al.*, 2014. *Sanchez 2017*: calibrations according to Sánchez-Baracaldo *et al.*, 2017. *GOE*: calibrations based on only the Great Oxidation Event. *Root-1*: a more conservative maximum age for the root.

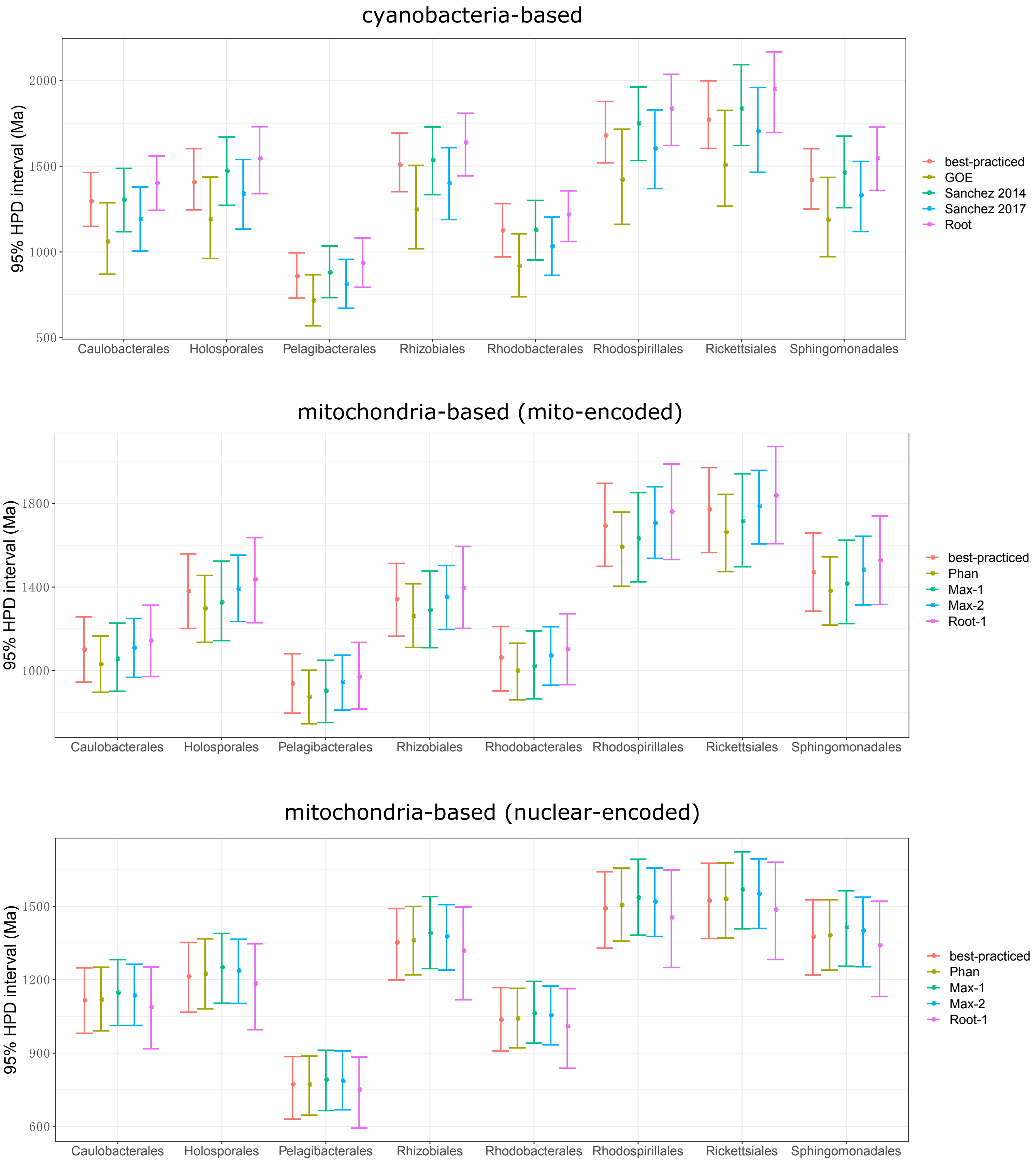

Figure S10. Selection of the best-fit clock model in MCMCTree analysis. (A) Results of mcmc3r analysis. The number of genes (g) and species (s) for each dataset are indicated. The tested clock models are the autocorrelated rates clock (AR), independent rates clock (IR), and the strict clock (STR). *Pr* indicates the posterior probabilities of different clock models, and those higher than 0.75 correspond to substantial evidence according to the evidence categories for the Bayes Factor given by Jeffreys, 1961. The model with the highest posterior probability (and above 0.75) is shown in bold type. The data for mcmc3r analysis are available at the online open access repository FigShare (see Data Availability). (B, C) Boxplots of the average substitution rates (amino acid changes per site per 100 million years) of branches belonging to the mitochondria, *Rickettsiales* and other alphaproteobacterial clades using the mito-encoded (B) and nuclear-encoded (C) dataset. The substitution rates were estimated by MCMCTree under the AR and IR model, respectively. The *P*-values were calculated using a paired t-test (* $p<0.05$, ** $p<0.01$, *** $p<0.001$, n.s.: not significant).

A

| | Mito-encoded | | | Nuclear-encoded | | |
|---|---|---|---|---|---|---|
| Data | Model | Log ML | Pr | Model | Log ML | Pr |
| 20s, 3g | AR | -29563.91 | 0.861 | AR | -24155.84 | 0.072 |
| | IR | -29565.73 | 0.139 | IR | -24153.29 | 0.928 |
| | STR | -29697.46 | 0 | STR | -24284.6 | 0 |
| 20s, 5g | AR | -53950.53 | 0.781 | AR | -37694.31 | 0.974 |
| | IR | -53951.8 | 0.219 | IR | -37697.95 | 0.026 |
| | STR | -54225.71 | 0 | STR | -37886.26 | 0 |
| 20s, 10g | AR | -91982.55 | 0.999 | AR | -97673.1 | 0.468 |
| | IR | -91989.26 | 0.001 | IR | -97672.97 | 0.532 |
| | STR | -92322.99 | 0 | STR | -98195.11 | 0 |
| 40s, 3g | AR | -35186.76 | 1 | AR | -32660.08 | 0.879 |
| | IR | -35196.25 | 0 | IR | -32662.06 | 0.121 |
| | STR | -35412.79 | 0 | STR | -32792.46 | 0 |
| 40s, 5g | AR | -81426.1 | 0.988 | AR | -65955.1 | 1 |
| | IR | -81430.48 | 0.012 | IR | -65966.05 | 0 |
| | STR | -81923.47 | 0 | STR | -66271.04 | 0 |
| 40s, 10g | AR | -156821.1 | 0 | AR | Not converged | |
| | IR | -156805.5 | 1 | IR | -126709.4 | N/A |
| | STR | -157618.8 | 0 | STR | -127333.6 | |

B

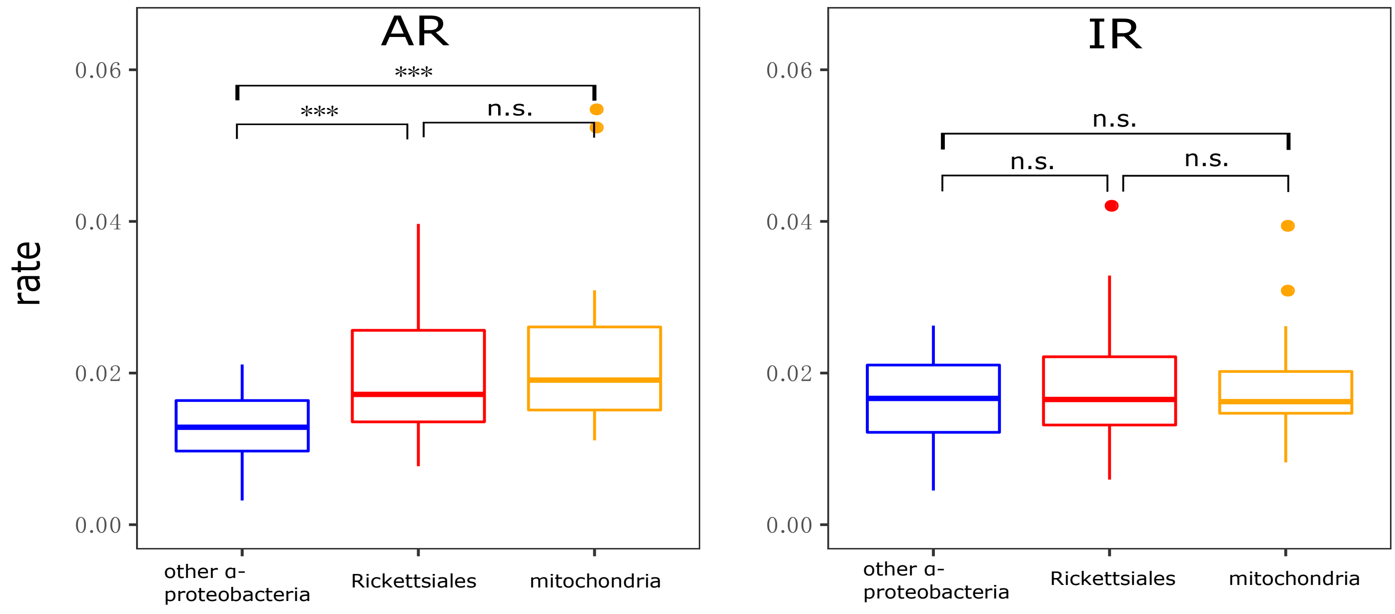


C

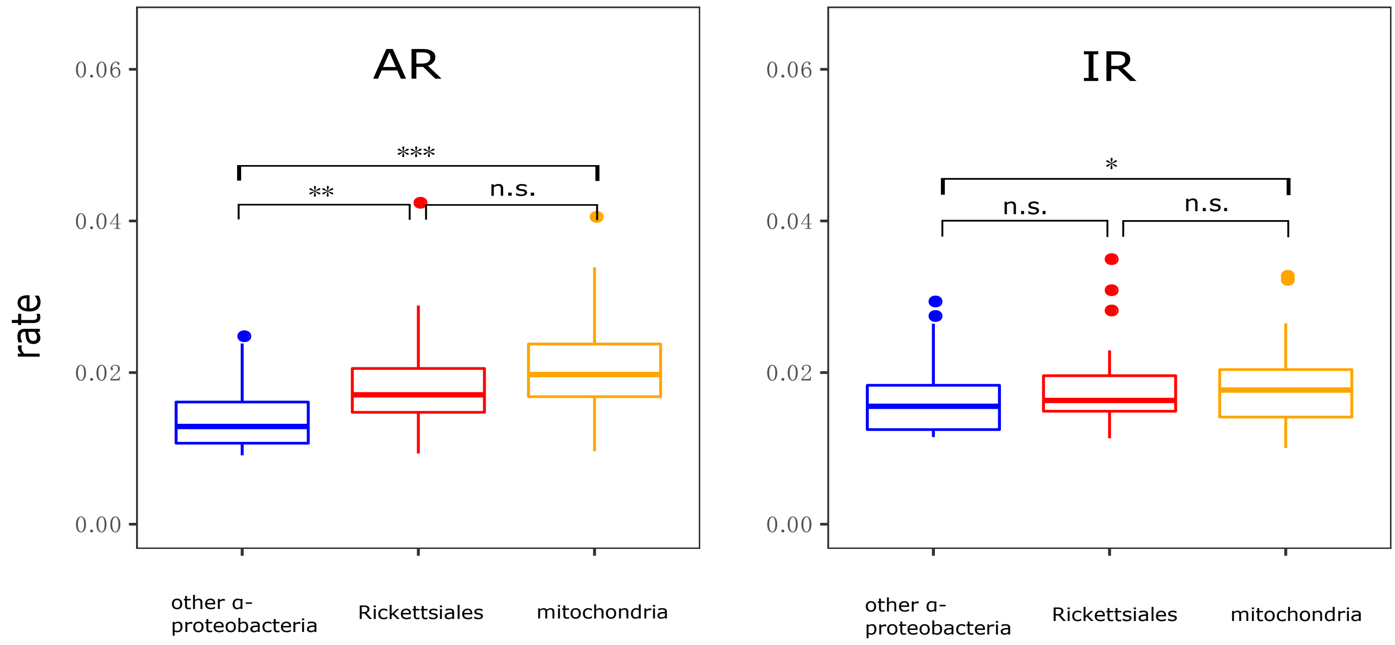

Figure S11. Reconstruction of ancestral hosts of *Rickettsiales* and transition rates between animal- and protist-associated lifestyles with randomly selected representatives from each OTU (defined by 98.7% identity of 16S rRNA gene). (A) Inferred ancestral hosts of *Rickettsiales*. The pie charts on the nodes show the estimated probabilities of the hosts, and the branch colors indicate the hosts with the higher probability at the corresponding node. (B) The transition rates from animal-associated to protist-associated ($q_{AP}$) and from protist-associated to animal-associated ($q_{PA}$) estimated by the MCMC method in BayesTraits multistate. The log-transformed Bayes factor (logBF) is indicated, where values above 10 are considered very strong evidence for support (Meade andPagel, 2016). The sources and credits of the cartoon graphs are provided in the online open access repository FigShare (see Data availability).

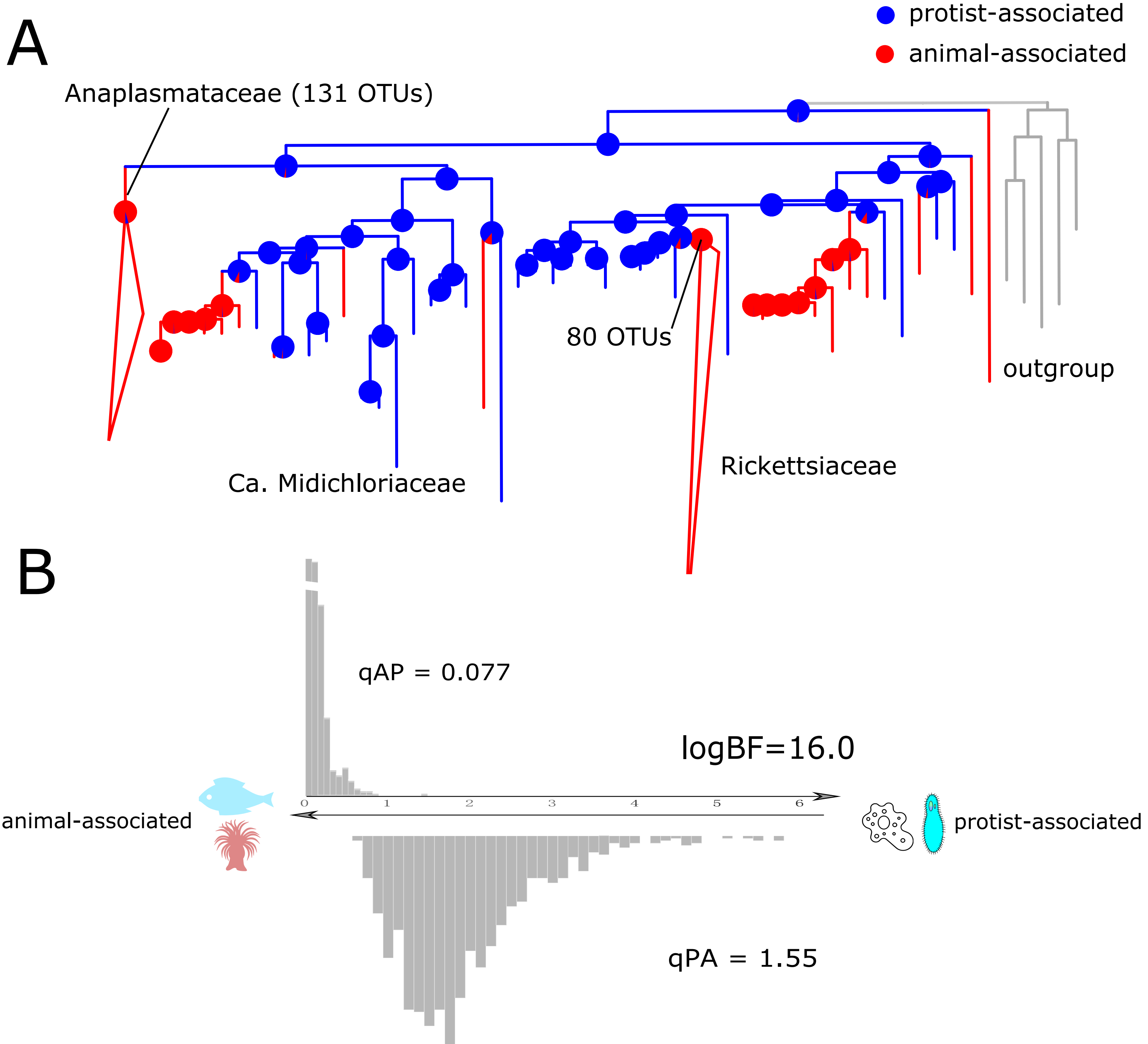

Figure S12.1 Visualization of the multiple sequence alignments of the five protein-coding genes that likely contain large proportion of ambiguous sites across bacteria, plastids and mitochondria analyzed in Shih *et al.*, 2017. The alignments were generated using MAFFT v7.222 with default settings. Amino acids are shaded according to the color scheme based on the *BLOCUM62* substitution matrix. (A) *AtpE*. (B) *AtpH*. (C) *AtpF*. (D) *Rpl2*. (E) *Rps3*.

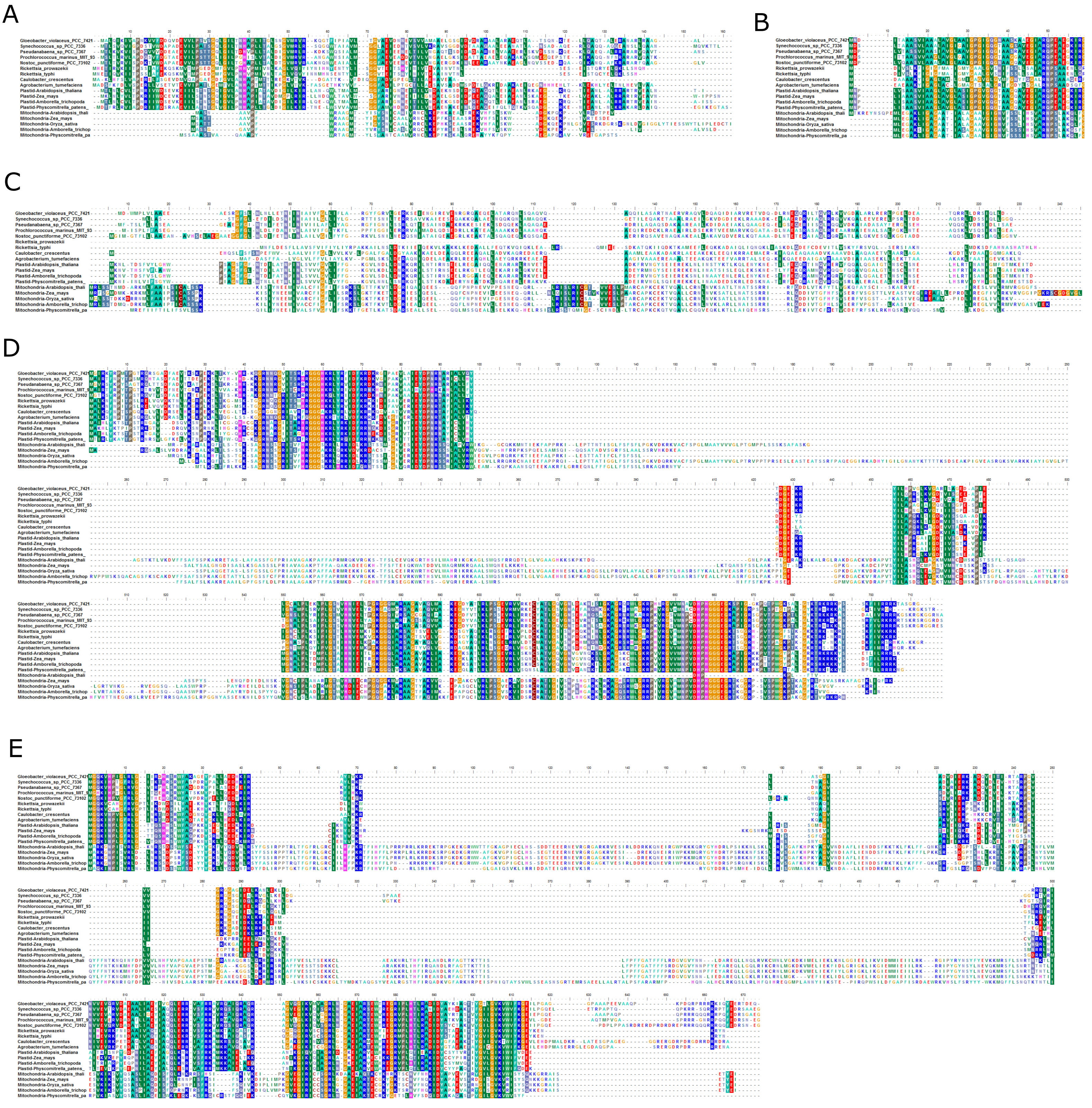

Figure S12.2 Visualization of the multiple sequence alignments of the other six protein-coding genes across bacteria, plastids and mitochondria analyzed in Shih *et al.*, 2017. The alignments were generated using MAFFT v7.222 with default settings. Amino acids are shaded according to the color scheme based on the *BLOCUM62* substitution matrix. (A) *AtpA*. (B) *AtpB*. (C) *AtpI*. (D) *EF-Tu*. (E) *Rpl16*. (F) *Rps12*.

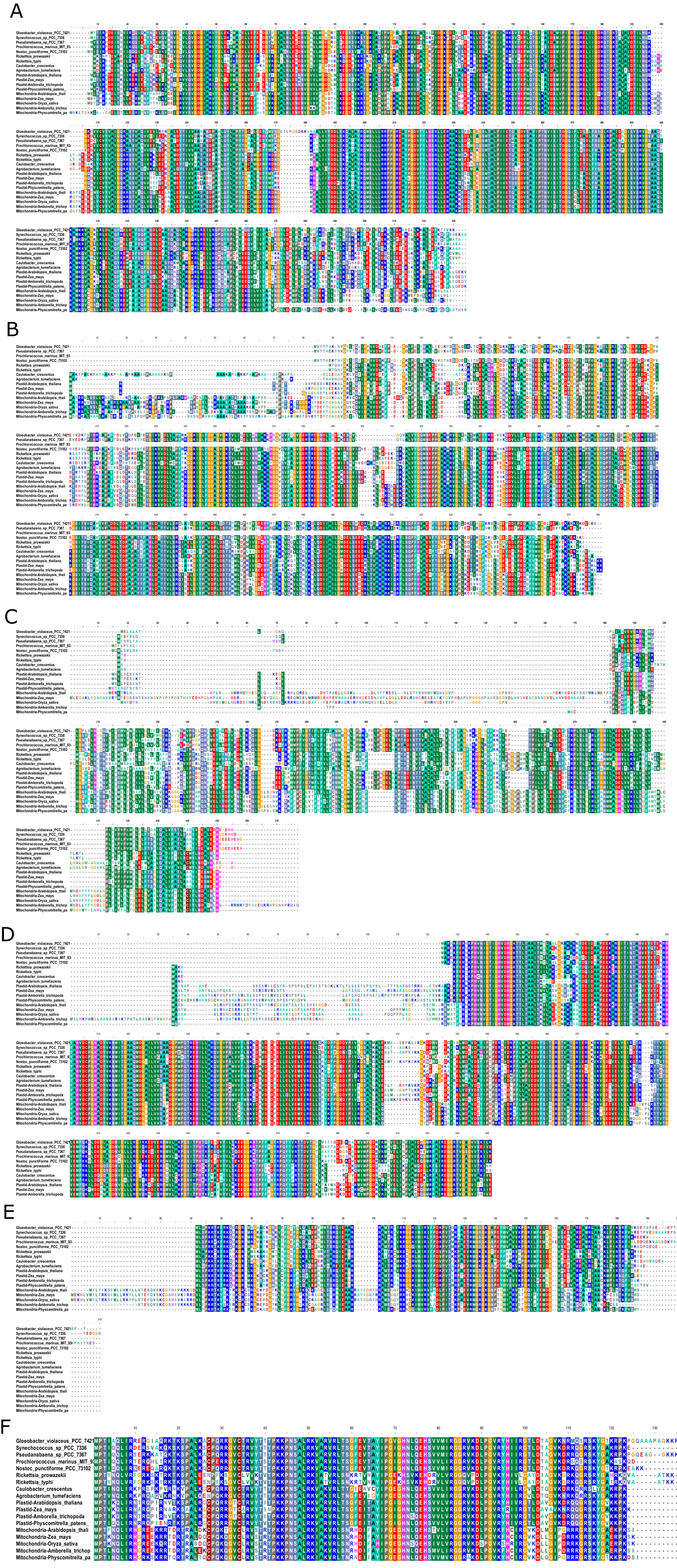
