## Supplementary material for "Dating *Alphaproteobacteria* evolution with eukaryotic fossils": Tables S1-S2, Notes S1-S3

**This file includes the following items**

Tables S1 to S2

Supplementary Notes S1 to S3

Supplementary references

**Table of contents**

Tables S1 to S2

Table S1 …………………………………………………………………………… 3

Table S2 …………………………………………………………………………… 7

Supplementary Notes S1 to S3

Note S1: Supplementary Methods

S1.1 Genome selection ……………………….………………………………. 8

S1.2 Molecular dating …………………..…………………………………….. 9

S1.3 Reconstruction of ancestral hosts ……………………..….....………………… 12

S1.4 Data visualization ………………..........................................……………… 12

Supplementary Note S2: Calibration information

Supplementary Note S3: Additional discussion

S3.1 Limitations of the approach adopted in Shih et al., 2017 …....………..…… 19

S3.2 Limitations of calibrating the evolution of bacteria based on the divergence times of their modern hosts ……………………………………………………….…….. 19

S3.3 Implications for the evolution of *Rickettsiales* …………………….……..… 20

S3.4 Selection of the clock model ……………………..………………………..… 22

Supplementary references

**Table S1.** Genome sources.

| Taxonomy | Species | Nuclear genome | Mitogenome |
|---|---|---|---|
| Metazoa | Homo sapiens | Ensembl release 96 (Howe et al., 2020) | |
| Metazoa | Gallus gallus | Ensembl release 96 (Howe et al., 2020) | |
| Metazoa | Branchiostoma floridae | UniProt (Bateman, 2019) | |
| Metazoa | Amphimedon queenslandica | Ensembl Metazoa release 46 (Howe et al., 2020) | |
| Fungi | Candida albicans | UniProt (Bateman, 2019) | |
| Fungi | Ustilago maydis | Ensembl Fungi release 46 (Howe et al., 2020) | |
| Fungi | Pleurotus ostreatus | UniProt (Bateman, 2019) | |
| Fungi | Spizellomyces punctatus | UniProt (Bateman, 2019) | |
| Amoebozoa | Acanthamoeba castellanii | Ensembl Protists release 43 (Howe et al., 2020) | |
| Amoebozoa | Dictyostelium discoideum | Ensembl Protists release 43 (Howe et al., 2020) | |
| Amoebozoa | Polysphondylium pallidum | dictyBase (Basu et al., 2013) | |
| Archaeplastida | Arabidopsis thaliana | PLAZA 4.0 (Van Bel et al., 2018) | MitoCOGs |
| Archaeplastida | Oryza sativa | PLAZA 4.0 (Van Bel et al., 2018) | MitoCOGs |
| Archaeplastida | Physcomitrella patens | PLAZA 4.0 (Van Bel et al., 2018) | MitoCOGs |
| Archaeplastida | Ostreococcus tauri | PLAZA 2.0 (Proost et al., 2009) | MitoCOGs |
| Archaeplastida | Chondrus crispus | Ensembl Plants release 46 (Howe et al., 2020) | MitoCOGs |
| Archaeplastida | Porphyra umbilicalis | UniProt (Bateman, 2019) | (Smith et al., 2012) |
| Archaeplastida | Cyanidioschyzon merolae | Ensembl Plants release 43 (Howe et al., 2020) | MitoCOGs |
| Archaeplastida | Cyanophora paradoxa | http://cyanophora.rutgers.edu/cyanophora/ | MitoCOGs |
| Heterokontophyta | Phytophthora infestans | Ensembl Protists release 46 (Howe et al., 2020) | MitoCOGs |
| Heterokontophyta | Thalassiosira pseudonana | Ensembl Protists release 46 (Howe et al., 2020) | |
| Discoba | Andalucia godoyi | https://megasun.bch.umontreal.ca/Andalucia_godoyi/ (Gray et al., 2020) | MitoCOGs |
| Discoba | Histiona aroides | | MitoCOGs |
| Discoba | Jakoba bahamiensis | | MitoCOGs |
| Discoba | Jakoba libera | | MitoCOGs |
| Discoba | Reclinomonas americana | | MitoCOGs |
| Discoba | Seculamonas ecuadoriensis | | MitoCOGs |
| Malawimonadida | Malawimonas jakobiformis | | MitoCOGs |
| SAR | Symbiodinium minutum | https://marinegenomics.oist.jp/ v1.2 (Shoguchi et al., 2013) | |
| SAR | Paramecium tetraurelia | Ensembl Protists release 45 (Howe et | |

| | | |
|---|---|---|
| | | al., 2020) |
| SAR | Oxytricha trifallax | OxyDB (http://oxy.ciliate.org/) |
| SAR | Reticulomyxa filosa | Ensembl Protists release 43 (Howe et al., 2020) |
| SAR | Elphidium margaritaceum | MMETSP (transcriptomes) (Keeling et al., 2014) |
| α-proteobacteria | Acidiphilium angustum ATCC 35903 | GCA_000701585.1 |
| α-proteobacteria | alpha proteobacterium AAP81b | GCA_001295935.1 |
| α-proteobacteria | alpha proteobacterium Q-1 | GCA_000710935.1 |
| α-proteobacteria | alpha proteobacterium RS24 | GCA_000469155.1 |
| α-proteobacteria | alphaproteobacterium SCGC AAA288-N07 | GCA_000513055.1 |
| α-proteobacteria | alphaproteobacterium sp. HIMB5 | GCA_000299095.1 |
| α-proteobacteria | Anaplasma phagocytophilum HZ | GCA_000013125.1 |
| α-proteobacteria | Asticcacaulis excentricus CB 48 | GCA_000175215.2 |
| α-proteobacteria | Bartonella quintana RM-11 | GCA_000294715.1 |
| α-proteobacteria | Caedibacter 37-49 | GCA_001898725.1 |
| α-proteobacteria | Caedibacter 38-128 | GCA_001898705.1 |
| α-proteobacteria | Caedibacter varicaedens | GCA_001192655.1 |
| α-proteobacteria | Candidatus Arcanobacter lacustris | GCA_000970895.1 |
| α-proteobacteria | Candidatus Caedibacter acanthamoebae | GCA_000743035.1 |
| α-proteobacteria | Candidatus Finniella lucida | GCA_004210305.1 |
| α-proteobacteria | Candidatus Jidaibacter acanthamoeba | GCA_000815465.1 |
| α-proteobacteria | Candidatus Midichloria mitochondrii IricVA | GCA_000219355.1 |
| α-proteobacteria | Candidatus Nucleicultrix amoebiphila FS5 | GCA_002117145.1 |
| α-proteobacteria | Candidatus Odyssella thessalonicensis L13 | GCA_000190415.2 |
| α-proteobacteria | Candidatus Paracaedibacter acanthamoebae isolate PRA3 | GCA_000742835.1 |
| α-proteobacteria | Candidatus Paracaedibacter symbiosus | GCA_000757605.1 |
| α-proteobacteria | Candidatus Pelagibacter IMCC9063 | GCA_000195085.1 |
| α-proteobacteria | Candidatus Pelagibacter ubique HTCC8051 | GCA_000472605.1 |
| α-proteobacteria | Citromicrobium JLT1363 | GCA_000186705.2 |
| α-proteobacteria | Ehrlichia canis Jake | GCA_000012565.1 |
| α-proteobacteria | Elioraea tepidiphila DSM 17972 | GCA_000378465.1 |
| α-proteobacteria | endosymbiont of Peranema | GCA_004210275.1 |
| α-proteobacteria | endosymbiont of Stachyamoeba | GCA_003932735.1 |
| α-proteobacteria | Granulibacter bethesdensis CGDNIH1 | GCA_000014285.2 |

| | | |
|---|---|---|
| α-proteobacteria | Hirschia baltica ATCC 49814 | GCA_000023785.1 |
| α-proteobacteria | Hyphomicrobium denitrificans ATCC 51888 | GCA_000143145.1 |
| α-proteobacteria | Inquilinus limosus DSM 16000 | GCA_000423185.1 |
| α-proteobacteria | Jannaschia EhC01 | GCA_001650845.1 |
| α-proteobacteria | Ketogulonicigenium vulgare WSH-001 | GCA_000223375.1 |
| α-proteobacteria | Kordiimonas gwangyangensis DSM 19435 - JCM 12864 | GCA_000375545.1 |
| α-proteobacteria | Meganema perideroedes DSM 15528 | GCA_000374145.1 |
| α-proteobacteria | Methylobacterium extorquens AM1 | GCA_000022685.1 |
| α-proteobacteria | Methylocystis SC2 | GCA_000304315.1 |
| α-proteobacteria | Micavibrio aeruginosavorus ARL-13 | GCA_000226315.1 |
| α-proteobacteria | Neorickettsia sennetsu Miyayama | GCA_000013165.1 |
| α-proteobacteria | Oceanibaculum indicum P24 | GCA_000299935.1 |
| α-proteobacteria | Oceanicaulis HTCC2633 | GCA_000152745.1 |
| α-proteobacteria | Orientia tsutsugamushi Boryong | GCA_000063545.1 |
| α-proteobacteria | Paracoccus denitrificans PD1222 | GCA_000203895.1 |
| α-proteobacteria | Parvibaculum lavamentivorans DS-1 | GCA_000017565.1 |
| α-proteobacteria | Parvularcula bermudensis HTCC2503 | GCA_000152825.2 |
| α-proteobacteria | Pelagibacter sp. HIMB058 | GCA_000012345.1 |
| α-proteobacteria | Pelagibacterium halotolerans B2 | GCA_000230555.1 |
| α-proteobacteria | Phenylobacterium zucineum HLK1 | GCA_000017265.1 |
| α-proteobacteria | Rhizobium leguminosarum bv. trifolii WSM1689 | GCA_000517605.1 |
| α-proteobacteria | Rhodobacter sphaeroides 2.4.1 | GCA_000012905.2 |
| α-proteobacteria | Rhodopseudomonas palustris TIE-1 | GCA_000020445.1 |
| α-proteobacteria | Rhodospirillum centenum SW | GCA_000016185.1 |
| α-proteobacteria | Rhodospirillum rubrum ATCC 11170 | GCA_000013085.1 |
| α-proteobacteria | Rhodovibrio salinarum DSM 9154 | GCA_000515255.1 |
| α-proteobacteria | Rickettsia typhi Wilmington | GCA_000008045.1 |
| α-proteobacteria | Rubritepida flocculans DSM 14296 | GCA_000425365.1 |
| α-proteobacteria | Ruegeria ANG-R | GCA_000813985.1 |
| α-proteobacteria | Sneathiella glossodoripedis JCM 23214 | GCA_000616095.1 |
| α-proteobacteria | Sphingomonas wittichii | GCA_000016765.1 |
| α-proteobacteria | Thalassobaculum salexigens DSM 19539 | GCA_000423805.1 |
| α-proteobacteria | Thalassospira profundimaris WP0211 | GCA_000300275.1 |
| α-proteobacteria | Wolbachia endosymbiont of Culex quinquefasciatus Pel | GCA_000073005.1 |
| α-proteobacteria | Wolbachia endosymbiont of Onchocerca ochengi | GCA_000306885.1 |
| Magnetococcales | Magnetococcus marinus MC-1 | GCA_000014865.1 |

| | | |
|---|---|---|
| Magnetococcales | Magnetofaba australis IT-1 | GCA_002109495.1 |
| β-proteobacteria | Burkholderia thailandensis E264 | GCA_000012365.1 |
| | Nitrosospira multiformis ATCC | GCA_000196355.1 |
| β-proteobacteria | 25196 | |
| γ-proteobacteria | Alteromonas lipolytica | GCA_001758465.1 |
| γ-proteobacteria | Spongiibacter tropicus DSM 19543 | GCA_000420325.1 |

**Table S2.** Comparison of the prior and posterior time estimates (mean and 95% HPD intervals) in MCMCTree analysis. The time estimates are given in the unit of million years.

**Mito-encoded**

| | **Prior** | | | **Posterior** | | |
|---|---|---|---|---|---|---|
| **Clade** | **Mean** | **Min** | **Max** | **Mean** | **Min** | **Max** |
| Mitochondria | 2333.06 | 1906.94 | 2715.27 | 1548.95 | 1709.19 | 1392.1 |
| Alphaproteobacteria | 2429.65 | 2124.34 | 2743.45 | 2036.73 | 1824.41 | 2265.25 |
| Caulobacterales | 1651.2 | 528.935 | 2456.39 | 1098.88 | 1257.58 | 944.449 |
| Holosporales | 1945.46 | 1134.97 | 2592.26 | 1379.05 | 1558.76 | 1201.44 |
| Rhizobiales | 1927.31 | 1183.52 | 2557.22 | 1340.98 | 1513.2 | 1164.4 |
| Rhodobacterales | 1708.09 | 671.579 | 2510.52 | 1061.49 | 1211.01 | 901.72 |
| Rhodospirillales | 2284.85 | 1840.1 | 2676.97 | 1692.5 | 1896.67 | 1499.16 |
| Rickettsiales | 2251.9 | 1711.28 | 2693.58 | 1770.91 | 1972.07 | 1565.49 |
| Pelagibacterales | 1840.16 | 813.189 | 2573.2 | 936.411 | 1080.31 | 796.013 |
| Sphingomonadales | 1902 | 915.604 | 2609.72 | 1470.26 | 1659.3 | 1284.38 |

**Nuclear-encoded**

| | **Prior** | | | **Posterior** | | |
|---|---|---|---|---|---|---|
| **Clade** | **Mean** | **Min** | **Max** | **Mean** | **Min** | **Max** |
| Mitochondria | 2368.22 | 1920.12 | 2798.79 | 1545.81 | 1429.56 | 1667.99 |
| Alphaproteobacteria | 2428.87 | 2063.87 | 2818.15 | 1771.92 | 1611.39 | 1931.68 |
| Caulobacterales | 1355.83 | 465.656 | 2456.05 | 1115.36 | 980.855 | 1248.74 |
| Holosporales | 1811.45 | 824.271 | 2667.37 | 1212.98 | 1067.23 | 1352.55 |
| Rhizobiales | 1722.34 | 706.049 | 2649.23 | 1351.88 | 1199.07 | 1491.05 |
| Rhodobacterales | 1423.3 | 499.92 | 2453.06 | 1035.79 | 908.287 | 1168.12 |
| Rhodospirillales | 2234.2 | 1606.03 | 2799.79 | 1393.35 | 1236.42 | 1546.76 |
| Rickettsiales | 2192.76 | 1435.45 | 2797.75 | 1327.27 | 1178.98 | 1475.16 |
| Pelagibacterales | 1618.46 | 576.052 | 2598.93 | 1522.6 | 1368.24 | 1676.63 |
| Sphingomonadales | 1698.44 | 576.559 | 2618.09 | 771.74 | 630.174 | 885.743 |

**Supplementary Note S1: Supplementary Methods**

**1.1 Genome selection**

Selection of genomes for phylogenomic reconstruction

We selected 80 genomes to determine the phylogenetic relationship of *Alphaproteobacteria* and mitochondria, including 64 alphaproteobacterial species, ten eukaryotes, and six outgroup genomes (Table S2). We chose 64 alphaproteobacterial genomes from a recent phylogenomic study of *Alphaproteobacteria* (Muñoz-Gómez et al., 2019) based on our following principles: i) Four fast-evolving genomes (*Candidatus* Hepatobacter penaei, *Holospora obtusa*, *Holospora undulata* and alphaproteobacterium HIMB59) and two genomes with unstable phylogenetic positions (*Tistrella mobilis* and *Geminicoccus roseus*) were removed; ii) All of the 12 and 9 genomes from *Rickettsiales* and *Holosporales*, respectively, were kept; iii) 43 genomes covering all major clades were selected to represent the diversity of *Alphaproteobacteria*. We chose ten mitochondrial genomes that are gene rich and that display relatively short branch lengths to minimize the impacts of long branch attraction, as used in a prior study (Martijn et al., 2018). These eukaryotic species were *Physcomitrella patens*, *Ostreococcus tauri, Andalucia godoyi*, *Jakoba libera*, *Jakoba bahamiensis*, *Histiona aroides*, *Seculamonas ecuadoriensis*, *Reclinomonas americana*, *Malawimonas jakobiformis*, and *Phytophthora infestans*. Six genomes from *Magnetococcales*, *Betaproteobacteria* and *Gammaproteobacteria* were used as the outgroup (Muñoz-Gómez et al., 2019).

Eukaryotic lineages used in dating

It is important to note that, as described above, we followed Martijn *et al.*, 2018 to construct the phylogeny of *Alphaproteobacteria* and mitochondria with ten eukaryotes whose mitogenomes are gene-rich and slowly-evolving thus ideal for phylogenetic analysis. However, these ten eukaryotes, which were mainly comprised by jakobids, a group of free-living heterotrophic protists from the eukaryotic supergroup Discoba, provided little fossil information. Hence, to take the advantages of eukaryotic fossils, we included more eukaryotic taxa in the mitochondria subtree for dating. We based the topology of the mitochondria subtree in dating analysis on the general consensus understanding of the eukaryotic phylogeny (Fig. S2A), rooted at the branch separating Amorphea (including Amoebozoa, Fungi and Metazoa) and others (Burki, 2014; Keeling and Burki, 2019). Using alternative topologies of the mitochondria subtree showed similar time estimates for *Alphaproteobacteria* (Fig. S8).

Eukaryotic lineages were selected based on prior studies (Betts et al., 2018; Martijn et al., 2018; Rodríguez-Ezpeleta and Embley, 2012) to cover both important lineages and those with high-quality fossil records. We compiled two datasets: the mito-encoded dataset, which was based on the 24 genes conserved between *Alphaproteobacteria* and mitochondrial genomes, and the nuclear-encoded dataset, which was based on 22 nuclear genes transferred from the mitochondrial genomes in evolution (Wang and Wu, 2015) (Dataset S1). Orthologs were retrieved from the MitoCOGs database and identified by BLAST search using sequences identified by Wang and Wu, 2015 as queries for the mito- and nuclear-encoded datasets, respectively, and only hits with the lowest e-values were retained (Table S1).

The genomes in the mito-encoded dataset included the ten aforementioned genomes used in phylogenomic reconstruction, and another six genomes from Archaeplastida (hereafter referred to as plants for simplicity) to take advantages of plant fossils (Fig. S2A), including two flowering plants (*Arabidopsis thaliana* and *Oryza sativa*), three red algae (*Chondrus crispus*, *Porphyra umbilicalis* and *Cyanidioschyzon merolae*), and a glaucophyte (*Cyanophora paradoxa*) (Fig. S2A). These genomes were chosen for the mito-encoded dataset because their mitogenomes were generally gene-rich and slowly-evolving (Martijn et al., 2018).

As shown in Fig. S2A, the nuclear-encoded dataset included all genomes used in the mito-encoded dataset except for the six without nuclear genomes, and additionally three amoebae, four animals, four fungi, and six from the Stramenopiles-Alveolata-Rhizaria (SAR) supergroup. Specifically, the three Amoebozoa lineages were two dictyostelids (social amoebae; *Dictyostelium discoideum* and *Polysphondylium pallidum*), and one Discosea species (flattened amoebae that move as a whole; *Acanthamoeba castellanii*). The four animals included two amniotes (*Homo sapiens* and *Gallus gallus*), a primitive chordate (*Branchiostoma floridae* [amphioxus]), and one from the early-split metazoan lineage sponge (*Amphimedon queenslandica*). The four fungi were three Dikarya species (*Candida albicans*, *Ustilago maydis* and *Pleurotus ostreatus*) and one chytrid, an early-split fungal lineage (*Spizellomyces punctatus*). The six SAR lineages consisted of a diatom (silicified microalgae; *Thalassiosira pseudonan*a), two rhizarians (a diverse group of protists defined by phylogenetic evidence; *Reticulomyxa filosa* and *Elphidium margaritaceum*), two ciliates [protists characterized by the presence of cilia; *Paramecium tetraurelia* from Oligohymenophorea and *Oxytricha trifallax* from Spirotrichea (Gao et al., 2016)], and a dinoflagellate (most members are characterized by two dissimilar flagella; *Symbiodinium minutum*). Including these eukaryotes allowed using six additional fossils and directly comparing the divergence times of host-associated bacteria and their eukaryotic hosts on the dated tree (Fig. S2A).

#### 1.2 Molecular dating

<u>MCMCTree analysis with the mitochondria-based strategy</u>

Previous studies showed that different partitioning schemes could lead to differences in the estimated ages and precision (Dos Reis et al., 2015; Foster and Ho, 2017). Genes within the same partition are allowed to have their own pattern of among-lineage rate heterogeneity in dating analyses. We considered a single partition (where all sequences were concatenated into one partition), the full partitioning (where each gene was regarded as an independent partition), and several other partitioning strategies (where genes with similar functions or features were clustered into the same partition). Different partitioning strategies showed generally consistent posterior estimates, in particular for the nuclear-encoded dataset (Fig. S6). In line with previous studies (Betts et al., 2018; Dos Reis et al., 2015), the fully partitioned strategy displayed the highest precision (Fig. S6), and was therefore used in the main analysis.

To determine the best-fit clock models, we employed the mcmc3r package (Dos Reis et al., 2018), which uses a stone-stepping method to calculate the marginal likelihood of three different clock models implemented in MCMCTree: the strict clock model (STR), the autocorrelated model (AR), and the independent rate (IR) model. This method works for the exact likelihood method, which is only available for nucleotide sequences in MCMCTree. To overcome the challenge imposed by the huge computational burden, we followed McGowen *et al.*, 2020: i) We reduced the subsets of taxa by randomly selecting 20 and 40 species for both the mito- and nuclear-encoded datasets; ii) For each subset, we randomly selected three, five and ten genes; iii) the root prior was fixed, and no other calibrations were used (here we did not intend to calculate divergence times but simply select the best-fit model). The lowest marginal likelihoods were always obtained with the strict clock model. The autocorrelated clock model turned out to have the highest marginal likelihood in most analyses (Fig. S10), and thus was used as the preferred model.

Phylogenetic studies have shown high rate heterogeneity among the mitochondria, *Rickettsiales* and the remaining lineages in *Alphaproteobacteria* (Fan et al., 2020; Martijn et al., 2018; Wang and Wu, 2015) (see also Fig. S1). We therefore set the parameter specifying the gamma distribution of the standard deviation of log-transformed rate on branches, σ, to be "1 10 1", which indicates a standard deviation of log-transformed rate on branches to be 0.1. This setting should accommodate considerable among-lineage rate variation (Brown and Yang, 2011). We also set σ to be "1 1 1" (meaning a standard deviation of log-transformed rate branches of 1.0) to accommodate larger among-lineage rate heterogeneity. The results were highly consistent (Fig. S5).

LG (Le and Gascuel, 2008) and GTR (Tavaré, 1986) were used as the substitution models for amino acid and nucleotide sequences, respectively. Four discrete categories of gamma distribution were applied to account for rate heterogeneity across sites. The mean substitution rate for each partition was based on the average substitution rates of all partitions calculated by codeml (or baseml for nucleotides) to inform the Dirichlet-gamma prior ("rgene_gamma").

MCMCTree analysis with the cyanobacteria-based strategy

Thirteen cyanobacteria genomes were selected based on a recent study (Wang et al., 2020) (Fig. S2B) after discarding three and six relatively closely related species from the *Pleurocapsales* and *Nostocales* respectively. Twenty-five conserved genes between *Alphaproteobacteria* and cyanobacteria used by others (Battistuzzi and Hedges, 2009; Wang et al., 2020) were analyzed. To keep consistency with the mitochondria-based analysis, the same settings in MCMCTree analysis were kept for the cyanobacteria-based method. Three calibration points within cyanobacteria were collected and used in the cyanobacteria-based dating analysis of *Alphaproteobacteria* (see Supplementary Note S2.2). The best-practiced calibration set was chosen based on careful review of the fossil evidence used in previous studies (see Supplementary Note S2.2).

Composite chronograms

The mean age of each node in the composite (joint) chronograms was calculated by integrating all the posterior ages obtained from different competing dating schemes (*Phan*, *Max-1*, *Root-1*, *Single partition*, *IR*, and best-practiced) using MCMCTree. The joint 95% HPD intervals were calculated using the R package BayesTwin (Schwabe, 2017). By integrating the estimates from different analyses, the composite chronograms can help better appreciate the uncertainties associated with Bayesian relaxed molecular clock analysis, as used in other studies (Betts et al., 2018; Dos Reis et al., 2015).

Infinite-sites plots

To assess whether the posterior density of the divergence times estimated by MCMCTree had reached the limiting distribution when the data had infinite sites, we employed the strategy employed by the previous studies (Rannala and Yang, 2007; Yang and Rannala, 2006). This theory predicts that given infinite amount of sequence data, the posterior mean of divergence times and the posterior credibility interval width should form a linear relationship, the slope in the infinite-sites plot denoting uncertainties in the estimated posterior ages that could not be reduced by sequence data alone. Hence, if in an infinite-sites plot the scatter points approach a straight line, the maximum amount of information has been obtained from the molecular data. This indicates that further reduction of the uncertainty in the posterior age estimates can be only achieved by using more informative calibration priors (Rannala and Yang, 2007).

Molecular dating using PhyloBayes

Molecular clock analysis of the divergence time of *Alphaproteobacteria* was also performed using PhyloBayes v4.1b (Lartillot et al., 2009) with both the log-normal autocorrelated ("-ln") rates and uncorrelated rates gamma multipliers ("-ugam") clock models, respectively. To keep consistency with the settings in the MCMCTree analysis, the root age was set as "-rp 2530 155", the LG model (Le and Gascuel, 2008) was used as the substitution model with four discrete gamma categories, and a soft tail of 2.5% to each of the upper and lower calibration bound was applied to all calibration points. Two independent chains were run, each with 60,000 iterations. The first 10,000 iterations were discarded as burn-in, and the rest was sampled every 10 iterations. Convergence was assessed using the "tracecomp" function implemented in PhyloBayes. The maximum bipartition discrepancies (maxdiff) across the two chains in all analyses were smaller than 0.1, indicating good runs (Lartillot et al., 2015).

Different topologies of the *Alphaproteobacteria* phylogeny

It is important to take into account the uncertainties associated with tree topology in divergence time estimation. Although our detailed phylogenomic reconstruction supports the topology shown in Fig. 1, there are several important unknowns about the phylogeny of the *Alphaproteobacteria*. The three main lineages that often displayed distinct phylogenetic positions in previous studies are mitochondria, *Holosporales* and *Pelagibacterales* (Roger et al., 2017). For these three lineages, we considered two, two and three different topologies, respectively, according to previous studies (Fig. S1B). The mitochondrial lineage was suggested to branch outside the *Alphaproteobacteria* (Martijn et al., 2018) (Topologies 1-3, 7-9), but are placed as a sister group of *Rickettsiales* according to traditional views (Andersson et al., 1998; Wang and Wu, 2015; Yang et al., 1985) (Topologies 4-6, 10-12). The *Holosporales* was originally thought to be closely related to *Rickettsiales* (Georgiades et al., 2011; Wang and Wu, 2015) (Topologies 7-12), but were shown to be closely related to *Rhodospirillales* when amino acid composition biases were corrected (Fan et al., 2020; Muñoz-Gómez et al., 2019) (Topologies 1-6). The *Pelagibacterales* was firstly identified as the sister group of mitochondria (Thrash et al., 2011) (Topologies 3, 6, 9, 12). However, later studies showed that their phylogenetic affiliation likely resulted from compositional biases, which, if accounted for, would result in a pattern where *Pelagibacterales* was within a clade with *Sphingomonadales*, *Rhizobiales*, *Caulobacterales* and *Rhodobacterales* (Topologies 2, 5, 8, 11) (Luo, 2015; Rodríguez-Ezpeleta and Embley, 2012). Consistent with recent studies with a more complete taxon sampling (Fan et al., 2020; Muñoz-Gómez et al., 2019), our phylogenomic analysis placed *Pelagibacterales* after its split with *Sphingomonadales* but prior to the divergence of *Rhizobiales*, *Caulobacterales* and *Rhodobacterales* (Topologies 1, 4, 7, 10).

#### 1.3 Ancestral lifestyle reconstruction of the *Rickettsiales*

A total of 2,321 16S rRNA gene sequences classified to *Rickettsiales* were downloaded from NCBI Genbank database (last accessed: June, 2020). These sequences were grouped into operational taxonomic units (OTUs) based on an identity cutoff of 97% (we also repeated the analysis with a more stringent cutoff of 98.7% [Fig. S11]) (Rodriguez-R et al., 2018). One sequence was randomly selected as the representative for each OTU. Ancestral hosts were inferred using the MCMC method from the multistate module in BayesTraits v3.0.2 (Meade and Pagel, 2016) based on the isolation hosts of extant members of *Rickettsiales*. The analysis was run for 1,100,000 iterations with the first 100,000 runs discarded as the burn-in. We calculated the marginal likelihoods with the stepping stone sampler implemented in BayesTraits with 100 stones each sampled for 1,000 iterations. The statistical significance of transition rates was assessed based on the log-transformed Bayes Factor (logBF) calculated as twice the difference of the log-transformed marginal likelihood between the two tested models (Meade and Pagel, 2016).

#### 1.4 Data visualization

Visualization of phylogenetic trees was performed using iTOL v4 (Letunic and Bork, 2019), FigTree v1.4.3 (http://tree.bio.ed.ac.uk/software/figtree), MEGA v7 (Kumar et al., 2016), TreeGraph v2.5 (Stöver and Müller, 2010), and the R package ggtree v2.0.4 (Yu et al., 2017). Multiple sequence alignments were visualized using BioEdit v7.0.5.3 (Hall, 1999).

**Supplementary Note S2: Calibration information**

Note that for all calibration points, both the minimum and maximum bounds are soft and there is a probability of 2.5% that the age is beyond the bound in our settings.

**2.1 Calibration points for the mitochondria-based strategy**

**Node:** Root (*Alphaproteobacteria* and *Beta-/Gammaproteobacteria* split)
**Datasets:** mito- and nuclear-encoded
**Minimum age:** 2220 Ma
**Maximum age:** 2840 Ma
**Justification:** We calibrated the prior of the root based on estimates from previous studies recorded in the database TimeTree (Kumar and Hedges, 2011). After removing duplicate records, there were seven time estimates for the split between *Alphaproteobacteria* and *Beta-/Gammaproteobacteria* (last accessed: March, 2020): 606 Ma (Chriki-Adeeb and Chriki, 2016), 2360 Ma (Sheridan et al., 2003), 2378 Ma (Battistuzzi and Hedges, 2009), 2504 Ma (Hedges and Kumar, 2009), 2508 Ma (Battistuzzi et al., 2004), 2621 Ma (Marin et al., 2017), and 2818 Ma (David and Alm, 2011). The estimate of 606 Ma differs dramatically from the others, and is likely flawed by their assumption of the concurrence of a couple of *Rhizobium* strains and their legume hosts in modern times (Wang et al., 2020). As a result, it probably underestimated the divergence time of all lineages in their phylogeny (see Supplementary Note S3.2 for more details). Thus, we discarded the estimate of 606 Ma, and derived a gamma-distributed prior of 2530 ± 153 Ma (2834-2226 Ma, 95% HPD interval) to the root based on the other six estimates.
**Alternatives**: We also used uniform prior distributions on the root age, which were 3000-2000 Ma, 3000-1500 Ma, and 3000-1000 Ma, corresponding to the dating scheme *Root-1*, *Root-2* and *Root-3* in Dataset S2, respectively. We used the gamma-distributed prior root age in the main analysis because i) it took less time to converge, ii) using uniformly-distributed root prior ages gave similar results, and iii) the gamma-distributed prior age of the root is compatible with the "-rp" setting in PhyloBayes.

**Node:** Crown group of Angiospermae (flowering plants) (Node 1 in Fig. S2A; both mito- and nuclear-encoded datasets)
**Locality and Stratigraphy level:** Cowleaze Chine Member, Isle of White
**Minimum Age:** 125 Ma
**Maximum Age:** 250 Ma
**Justification:** Tricopolate pollen is the most ancient evidence of angiosperms. Following Clark et al., 2011, we set the pollen to the Cowleaze Chine Member of the Vectis Formation, corresponding to a minimum time of 126.3 ± 0.4 Ma. The soft maximum time constraint was derived from sediments devoid of angiosperm-like pollen below their first report in the Middle Triassic, corresponding to 247.1 Ma ± 0.2 Ma (Ogg, 2012).

**Node:** Crown group of Embryophyta (land plants) (Node 2 in Fig. S2A; both mito- and nuclear-encoded datasets)
**Locality and Stratigraphy level:** Qusaiba-1 core from the Quasim formation of northern Saudi Arabia

**Minimum Age:** 450 Ma
**Maximum Age:** 509 Ma
**Justification:** Trilete spores are the oldest evidence of embryophytes known to date. We followed Clarke *et al.*, 2011 and dated these to 450 Ma, representing the minimum age of the crown group of embryophytes. The soft maximum constraint was placed at the Bright Angel Shale of the Tonto Group of Arizona with an age of 507.2-509 Ma (Baldwin et al., 2004), thus, the upper limit to 509 Ma (Clark and Donoghue, 2017).
**Alternatives:** As some molecular dating studies argued for an earlier origin of land plants (Hedges et al., 2018, 2004), we alternatively removed the maximum constraint of the age of this node in the dating scheme *Max-1* (Dataset S2).

**Node:** Total group of Florideophyceae (Node 3 in Fig. S2A; both mito- and nuclear-encoded datasets)
**Locality and Stratigraphy level:** Doushantuo Formation, southern China
**Minimum Age:** 550 Ma
**Maximum Age:** -
**Justification:** The anatomically preserved florideophyte fossils found at Doushantuo Formation displayed features that resemble reproductive structures of modern corallines (Xiao et al., 2004). Stratigraphically constrained by Nantuo glaciation and Ediacaran fossils, the age of Doushantuo fossils were estimated to be between 600 and 550 Ma (Knoll and Xiao, 1999). We followed Parfrey *et al.*, 2011, and used 550 Ma as the minimum age of the total group of Florideophyceae.

**Node:** Crown group of Rhodophyta (red algae) (Node 4 in Fig. S2A; both mito- and nuclear-encoded datasets)
**Locality and Stratigraphy level:** Lower Hunting Formation, Somerset Island, arctic Canada
**Minimum Age:** 1033 Ma
**Maximum Age:** -
**Justification:** The fossils of *Bangiomorpha pubescens* represent the oldest fossil that can be confidently assigned to a major eukaryotic clade. Based on the simple multicellular fossils with reproductive cell and differentiated holdfasts, people described *Bangiomorpha pubescens* as a Bangiales red algae (Knoll, 2011), and used it as the calibration point of the red algae crown group in other studies (Parfrey et al., 2011; Yang et al., 2016). We constrained the minimum age based on the date of a shale layer in the Artic Bay formation, which is 1092 ± 59 Mya (Turner and Kamber, 2012).
**Alternatives:** Some argued that the features considered to be characteristic of *Bangiomorpha* might also be found in other red algae (Betts et al., 2018). Thus, we followed Betts *et al.*, 2018 to assign the minimum age of the total group (Node a in Fig. S2A), instead of the crown group, of red algae to be 1033 Ma (scheme *Max-1* in Dataset S2).

**Node:** Crown group of Foraminifera (Node 5 in Fig. S2A; nuclear-encoded dataset)
**Locality and Stratigraphy level:** The Chapel Island Formation, Newfoundland, Canada
Minimum age: 525 Ma

Maximum age: -
**Justification:** The fossils of *Platysolenites cooperi* are considered the oldest one in Foraminifera. The wall composition indicates that *P. cooperi* is an agglutinating foraminifera (McIlroy et al., 2001). *P. cooperi* was discovered in the latest Ediacaran to Lower Cambrian in Newfoundland, the Chapel Island formation. According to the latest geological timescale (Gradstein et al., 2012), a minimum constraint of 525.5 Ma was set to this formation, thus the crown group of the Foraminifera.

**Node:** Crown group of Amniota (mammals, birds, and reptiles) (Node 6 in Fig. S2A; nuclear-encoded dataset)
**Locality and Stratigraphy level:** Joggins Formation of Nova Scotia, Canada
**Minimum age:** 312 Ma
**Maximum age:** 332 Ma
**Justification:** The oldest fossils that are assigned to Amniota are those of *Hylonomus lyelli* Dawson. Its minimum age was set to be 318 Ma based on the date of the Joggins Formation. We followed the study (Benton et al., 2015), and assigned the maximum age to be 332.9 Ma, the same as the fossiliferous Little Cliff Shale of the East Kirkton locality, where no fossils of reptilians are detected.
**Alternatives**: Despite little controversy in the maximum age of this node, to be conservative and to keep consistency with other nodes within the Metazoa, we removed the maximum age of this node in the dating scheme *Max-1* (Dataset S2).

**Node:** Crown group of Chordata (Node 7 in Fig. S2A; nuclear-encoded dataset)
**Locality and Stratigraphy level:** Haikou [Yuanshan Fm (formerly Qiongzhusi)], China
**Minimum Age:** 520 Ma
**Maximum Age:** 636 Ma
**Justification:** We followed Benton *et al.*, 2015 to set the minimum and maximum time constraints of the Chordata crown group. The minimum date was based on the fossils of *Haikouichthys ercaicunensis* (Shu et al., 1999), which were found in the Chengjiang Biota within the Yu'anshan Member of the Heilinpu Formation (Hou et al., 2008). There are many Lagerstätten (sedimentary deposits displaying extraordinary fossils with exceptional preservation) in the Lantian Biota preserving the biota, but none of these fossils shows any characteristics that could assign them to the Chordata crown group or even the eumetazoan total group (Benton et al., 2015). Thus, the appearance time of Chordata should not be earlier than the maximum age of the Lantian Biota, which is 636 Ma (Yuan et al., 2011).
**Alternatives**: As some argued for a more ancient origin of Chordata (Blair and Hedges, 2005), we removed the maximum age for this node in the dating scheme *Max-1* (Dataset S2).

**Node:** Crown group of Metazoa (animals) (Node 8 in Fig. S2A; nuclear-encoded dataset)
**Locality and Stratigraphy level:** White Sea Formation, Russia
**Minimum Age:** 550 Ma
**Maximum Age:** 833 Ma
**Justification:** The fossil of *Kimberella quadrata*, which belongs to Bilateria, is considered as the oldest metazoan's fossil. A lower limit of 550 Ma was established for *Kimberella*

*quadrata* (Fedonkin et al., 2007), and thus was used as the minimum age of Metazoa. A maximum age was established from the Svanbergfjellet Formation of Spitsbergen (Butterfield et al., 1994) and the Bitter Springs Formation of central Australia (Schopf, 1968) which preserve a variety of fossils assigned to eukaryotes (e.g., multicellular algae, sphaeromorph acritarchs) but show no evidence of total group metazoans. The absolute age of the Bitter Springs Formation was dated at 827 ± 6 Mya, therefore we used 833 Ma as the maximum constraint of Metazoa, as used in the previous studies (Betts et al., 2018; Dos Reis et al., 2015). Some studies based the minimum time of the animal crown group on the fossil lipids 24-isopropylcholestane and 26-methylstigmastane, dating back to 630 Mya. The reason was that these biomarkers were thought to be only associated with sponges (Parfrey et al., 2011), the presumably earliest-branching animal lineage (Feuda et al., 2017; Pisani et al., 2015). However, these biomarkers were later found to be common in rhizaria, an early-branching lineage of eukaryotes, hence the affinities of these biomarkers are ambiguous (Nettersheim et al., 2019).
**Alternatives**: According to some molecular dating studies (Cartwright and Collins, 2007; Hedges et al., 2004), animals might have originated 300-500 Ma earlier than the above estimate. Therefore, we alternatively removed the maximum constraint for this node (scheme *Max-1* in Dataset S2).

**Node:** Total group of Fungi (or crown group of Opisthokonta; Node 9 in Fig. S2A; nuclear-encoded dataset)
**Locality and Stratigraphy level:** Brock Inlierm, the Northwest Territories, Canada
**Minimum age:** 890 Ma
**Maximum age:** -
**Justification:** The oldest fungal fossil came from the shale of Grassy Bay Formation (Shaler Supergroup, Arctic Canada), described as *Ourasphaira giraldae*. It dates to 1010–890 Mya. *O. giraldae* shows septate hyphae that are characteristic of the Dikarya but that are also found in a few lineages of the Zoopagomycota, Mucoromycota, Chytridiomycota and Blastocladiomycota (Loron et al., 2019). Distinct from Dikarya, in the microfossil specimens of *O. giraldae*, researchers did not detect septa that are regularly distributed along the hyphae (Loron et al., 2019). These suggest that *O. giraldae* either is in a clade of the total group of fungi or belongs to the stem group of Dikarya (Loron et al., 2019). The oldest report of a fossil with features typical to fungi is found in the Ongeluk Formation, which dates back to 2400 Mya (Bengtson et al., 2017). However, this was questioned by Berbee *et al.*, 2017 because the morphology and deep marine habitat of the fossils were inconsistent with predictions from phylogenetic analysis. Further, its biological activities were thought to be suspicious as it lacks organic components (Berbee et al., 2017). To be conservative, we assigned a minimum time constraint of 890 Mya to the total group of fungi, i.e., the crown group of Opisthokonta.

**Node:** Crown group of Dikarya (Ascomycota and Basidiomycota) (Node 10 in Fig. S2A; nuclear-encoded dataset)
**Locality and Stratigraphy level:** Rhynie, Aberdeenshire, Scotland, Lower Devonian
**Minimum age:** 400 Ma

**Maximum age:** -
**Justification:** The earliest uncontroversial fossil that belongs to Dikarya is the fossil of *Paleopyrenomycites devonicus*, which shows key characteristics related to Ascomycota. The estimated date of the fossil was based on that of the Rhynie Chert system, which is ~400 Ma (Mark et al., 2011; Schoene et al., 2010).

### 2.2 Calibration points for the cyanobacteria-based strategy

**Node:** Root (cyanobacteria-alphaproteobacteria split)
**Minimum age:** 2320 Ma
**Maximum age:** 3500 Ma
**Justification:** The minimum constraint of the root age was based on geochemical estimate of the Great Oxidation Event [GOE; a time period when the oxygen concentrations increased by several orders of magnitude in the Earth's atmosphere (Eguchi et al., 2020; Kump, 2008)], which is often considered as a result of oxygenic photosynthesis by cyanobacteria (Kopp et al., 2005; Schirrmeister et al., 2013; Van Kranendonk et al., 2012). The maximum constraint was more difficult to determine because modern phylogenetics analysis showed that cyanobacteria likely have a very deep phylogenetic position whereas fossils of early life are often under controversy (Javaux, 2019). We chose 3500 Ma as the maximum constraint since better-preserved fossils indicating activities of early life were dated back to ~3500 Myr (Allwood et al., 2007; Javaux, 2019; Strauss, 1993; Van Kranendonk et al., 2008).
**Alternatives:** An alternative maximum age for the root of 3800 Ma (Nisbet and Sleep, 2001; Nutman et al., 2016) was used as some suggest that early life, thus the divergence between cyanobacteria and alphaproteobacteria, could occur earlier (dating scheme *Root-1* in Dataset S2).

**Node:** *Pleurocapsales* crown group (Node 1 in Fig. S2B)
**Minimum age:** 1700 Ma
**Maximum age:** 1900 Ma
**Justification:** For the minimum age, it was based on the estimated age of the microfossils of *Pleurocapsales* found in Hebei, China, which dates back to ca. 1700 Mya (Zhang and Golubic, 1987). The setting of the soft maximum constraint was according to the studies (Golubic and Seong-Joo, 1999; Sergeev et al., 2002) which suggested that microfossils with sheaths and increased cell diameters, which are typically characteristic to *Nostocales* and *Pleurocapsales*, became popular after 1900 Mya. The combination of minimum and maximum ages were also used in previous studies (Sánchez-Baracaldo, 2015; Sánchez-Baracaldo et al., 2017).
**Alternatives:** It was inferred that the ancestor of *Pleurocapsales* had cell diameters of large size, which, according to some studies, did not arise until 2450 Mya (Blank and SÁnchez-Baracaldo, 2010; Tomitani et al., 2006). We therefore used this time information as an alternative maximum constraint for the age of the *Pleurocapsales* (Sánchez-Baracaldo et al., 2014) (dating scheme *Sanchez 2014* in Dataset S2).

**Node:** *Nostocales* crown group (Node 2 in Fig. S2B)

**Minimum age:** 1200 Ma
**Maximum age:** 2100 Ma
**Justification:** There are several fossils described as akinetes, which are considered to be characteristic to the *Nostocales*. Some of the very ancient records are the ones preserved in Franceville Group of Gabon (Amard and Bertrand-Sarfati, 1997) and McArthur Group of Northern Australia (Tomitani et al., 2006), dating back to ~2100 Ma and ~1600 Ma, respectively. However, their validity was questioned (Butterfield, 2015). Thus, we opted for a more conservative minimum age of this node based on a younger fossil (1.2 Ga) (Horodyski and Allan Donaldson, 1980) with better morphological evidence (Butterfield, 2015). For the maximum age, it was suggested that the heterocyst of *Nostocales* originated as a response to oxygen. The oxygen in the atmosphere might not reach the levels allowing for the differentiation of heterocyst at ~2100 Mya (Tomitani et al., 2006). Thus, we used this time information for the upper time boundary of this node as used in other studies (Sánchez-Baracaldo, 2015).
**Alternatives:** We also tried two older estimates of the akinete fossils mentioned above, which were 2450-2100 Ma (Sánchez-Baracaldo et al., 2014) and 1900-1600 Ma (Sánchez-Baracaldo et al., 2017) (schemes *Sanchez 2015* and *Sanchez 2017* in Dataset S2).

**Node:** cyanobacteria crown group (Node 3 in Fig. S2B)
**Minimum age:** 2320 Ma
**Maximum age:** 3000 Ma
**Justification:** The lower limit was set based on the time estimate of GOE (see above). The upper limit was on the basis of sensitive redox proxies which suggested an earlier origin of cyanobacteria dating back to around 3000 Ma (Crowe et al., 2013), as also used in other studies (Sánchez-Baracaldo, 2015; Sánchez-Baracaldo et al., 2017, 2014).

### Supplementary Note S3: Additional discussion

#### S3.1. Limitations of the approach adopted in Shih *et al.*, 2017

Shih *et al.*, 2017 pioneered using the idea of endosymbiosis to date the evolution of cyanobacteria by analyzing 12 homologs shared by cyanobacteria, plastids, and mitochondria. However, since there are some cyanobacteria fossils well suited for molecular dating (Sánchez-Baracaldo et al., 2017; Schirrmeister et al., 2016), it is unclear how much it improves the dating by integrating eukaryotic fossils. More importantly, mitochondria and plastids diverged long time ago, and both have undergone extensive gene reduction (de Vries and Archibald, 2018; Wang and Wu, 2015), which led to relatively few shared high-confidence orthologs. We found that five out of the 12 genes used by Shih *et al.*, 2017 were likely to contain large proportions of ambiguous sites in alignment (Fig. S12), which could result in overestimates of the substitution rates of these genes, thus underestimates of divergence time. Possibly because of this issue, the origin times of the crown group of cyanobacteria estimated by Shih *et al.*, 2017 were at least 400 Ma younger than estimated by others (Sánchez-Baracaldo, 2015; Sánchez-Baracaldo et al., 2017; Schirrmeister et al., 2013). Particularly, their estimated origin time of *Nostocale*s (~500 Ma) was much younger than the fossil records [before 1200 Ma (Horodyski and Allan Donaldson, 1980; Schirrmeister et al., 2016)].

#### S3.2. Limitations of calibrating the evolution of bacteria based on the divergence times of their modern hosts

Our time estimates of the divergences of *Alphaproteobacteria* were broadly consistent with the previous studies (Battistuzzi et al., 2004; Battistuzzi and Hedges, 2009), but were considerably older than others (Chriki-Adeeb and Chriki, 2016; Luo et al., 2013; Weinert et al., 2009). The first two studies (Battistuzzi et al., 2004; Battistuzzi and Hedges, 2009) calibrated the bacterial tree of life based on cyanobacteria fossils and/or other geologic records and biomarkers. In contrast, all of the latter three studies (Chriki-Adeeb and Chriki, 2016; Luo et al., 2013; Weinert et al., 2009) based calibration bound constraints (in part) on a strict host-bacteria co-evolution. In other words, they assumed that pathogenic/symbiotic alphaproteobacterial lineages co-diverged with their modern hosts, and used the fossils of their modern hosts to calibrate the evolution of *Alphaproteobacteria*. However, none of them considered the possibility of host shifts in evolution, which, if indeed occurring, can lead to underestimates of the divergence times of bacteria.

Specifically, Chriki-Adeeb and Chriki, 2016 based their analysis on a single calibration. They assumed that *Rhizobium sullae* originated at the same time of their modern hosts, i.e., the Hedysaroid clade (Fabaceae: Hedysareae), 29.3 ± 3.0 Mya. However, host transitions within legumes are very frequent for rhizobia (Masson-Boivin et al., 2009; Mutch and Young, 2004; Wang et al., 2012; Young and Johnston, 1989). It is possible that the LCA of *R. sullae* established a symbiosis relationship with other legumes and had its hosts shifted later in evolution. This means that the symbiosis between *R. sullae* and Hedysaroid might have started after the divergence of *R. sullae*. If so, the use of the divergence time of the legume hosts (Hedysaroid) as the occurrence time of the LCA of *R. sullae* could lead to an underestimate of the origin time of *R. sullae* and of all analyzed alphaproteobacteria, as pointed out by Wang *et al.*, 2020. Consequently, the divergence time between

*Alphaproteobacteria* and *Betaproteobacteria* estimated by Chriki-Adeeb and Chriki, 2016 was only 600 Ma.

Luo *et al.*, 2013 used three calibration points, two of which were based on fossil or geochemical evidence from cyanobacteria. As to the other one, it was based on the interaction between rhizobia and legumes, which was originally derived from an earlier study (Ochman and Wilson, 1987). Assuming that *Rhizobium* diverged from their non-rhizobia relatives after the appearance of legumes, Luo *et al.*, 2013 constrained the split of *Rhizobium* and its close relative (*Agrobacterium*) to be 120-100 Ma, the estimated origin time of legumes. This could be true if the LCA of *Rhizobium* indeed nodulated legumes. However, recent genomics studies showed that many *Rhizobium* strains, particularly the early-split lineages, do not adapt to a nodulating lifestyle, but are instead associated with diverse non-legume plants (Garrido-Oter et al., 2018; Wang et al., 2020; Yeoh et al., 2017). Hence, the origin time of legumes is plausibly more recent than the *Rhizobium-Agrobacterium* split. As a result, the assumption made in Luo *et al.*, 2013 (and originally in Ochman and Wilson, 1987) likely led to underestimates of the divergence times in the phylogeny.

Weinert *et al.*, 2009 dated the origin time of *Rickettsiales* to be 525-425 Ma. This study was based on the substitution rate of 16S rRNA. The substitution rate estimate was originally derived from an earlier study (Moran et al., 1993), where the authors assumed a co-divergence between *Buchnera* and its host (viz. aphids), and assigned the origin time of aphids as that of *Buchenera*. Similar to the study Chriki-Adeeb and Chriki, 2016, Weinert *et al.*, 2009 did not take into consideration the possibility of host shifts in evolution. Further, the substitution rate of the gene in *Buchnera*, which belongs to the *Gammaproteobacteria*, could be very different from that in *Rickettsiales*. Moreover, Weinert *et al.*, 2009 assumed a strict clock across the entire phylogeny. In other words, the authors assumed that non-*Rickettsiales* lineages, most of which are not parasitic bacteria, evolved at the same rate as *Rickettsiales*. Thus, this likely resulted in an overestimate of the substitution rate of non-*Rickettsiales* lineages, hence an underestimate of the divergence time between *Rickettsiales* and other alphaproteobacterial lineages.

#### S3.3. Implications for the evolution of *Rickettsiales*

People's understanding of *Rickettsiales* had long been limited to medically relevant members from the genera *Anaplasma*, *Ehrlichia*, *Neorickettsia* and *Rickettsia* (Thomas et al., 2016), and to *Wolbachia*, which infect 20-75% of insect species and almost all filarial nematodes (reviewed in Bordenstein, 2003; Werren *et al.*, 2008). Thus, the predominant view, proposed by Weinert *et al.*, 2009, is that the LCA of *Rickettsiales* already had the ability to infect arthropods and shifted to other eukaryotes later ("animal first"). More recently, accumulating evidence indicates a much broader range of hosts of *Rickettsiales*, which expanded to annelids, sponges, cnidarians (Klinges et al., 2019; Perlman et al., 2006), green algae (Kawafune et al., 2014, 2012), and diverse protists particularly amoebae and ciliates (Montagna et al., 2013; Szokoli et al., 2016; Vannini et al., 2005). An intriguing alternative hypothesis is therefore a "protist first" view where early *Rickettsiales* were associated with protists and switched to animals during evolution (Schrallhammer et al., 2013).

As detailed in Supplementary Note S3.1, the dated ages (525-425 Ma) of the crown group of *Rickettsiales* by Weinert *et al.*, 2009 is likely to be largely underestimated due to methodological biases. This study has been widely cited by others (Baldridge et al., 2010; De La Fuente et al., 2015; El Karkouri et al., 2016; Krawczak et al., 2018; Merhej and Raoult, 2011; Sachman-Ruiz and Quiroz-Castañeda, 2018; Tomassone et al., 2018), and might have a large impact on the field. Our revisited evolutionary timeline indicates that the LCA of *Rickettsiales* originated 1500 Mya, much earlier than the origin of animals but coincided with that of (unicellular) eukaryotes. Our results reject the view of the concurrence of the LCA of *Rickettsiales* and animals, and provide strong support to the "protist first" hypothesis. The reason is straightforward: if the parasites arose before the origin of their modern hosts, their interaction must be established later in parasites' evolution. The dating result is reinforced by ancestral host reconstruction, which inferred a protist-associated LCA of *Rickettsiales* with a posterior probability of 96.5% and a substantially higher host transition rate from animals to protists compared to that from protists to animals (Fig. 2D-E). Clearly, this points to independent host shifts from unicellular eukaryotes to animals, which, according to Fig. 2D, occurred once in the LCA of *Anaplasmataceae* whose all extant members are animal parasites, four times within the *Rickettsiaceae* where most members are associated with animal species, and three times in *Ca.* Midichloriaceae which are mostly associated with protists. Considering the diversity of *Rickettsiales* members yet to be studied and because the 16S rRNA genes used in our ancestral host reconstruction were only from representatives of each OTU, such evolutionary shifts are plausibly more frequent than here estimated. Note that a recent study described a new *Rickettsiales* family *Deianiraea*, which were reported to be attached to the surface of the ciliate *Paramecium* but never invading the inside of the host cell (Castelli et al., 2019). This opens a possibility that the LCA of *Rickettsiales* adapted to an extracellular parasitic lifestyle or was a facultatively intracellular bacterium. Nevertheless, it is clear that interaction between protists and *Rickettsiales* has a prominent role in *Rickettsiales* evolution and diversification.

*Rickettsiales* inhabiting hematophagous arthropods are transmitted to humans by either the bite of infected mites and ticks or the feces of infected lice and fleas that are inhaled or rubbed into the skin (Walker, 1996), which, however, cannot be the case for those inhabiting protists. It is hypothesized that protist-harboring *Rickettsiales* could be transmitted to animals sharing the same habitat via the feeding behaviour of the animal host, which presumably happened in aquatic environments where a variety of protists share the same habitats with invertebrates with grazing- or filter-feeding strategies (Kang et al., 2014; Schrallhammer et al., 2013). This view is further supported by the recent finding of the transfer of a *Rickettsiales* bacterium (*Ca.* Trichorickettsia mobilis) to a metazoan (the planarian *Dugesia japonica*) from an infected ciliate *Paramecium* in the same aquatic environment (Modeo et al., 2020). The common protist hosts of *Rickettsiales* such as amoebae or ciliates feed on bacteria, and are capable of harboring stable and rich prokaryotic communities, meaning that they can repeatedly take up bacteria from surrounding environments (Boscaro et al., 2019; Szokoli et al., 2016). As a feedback, protists provide protection against stresses and opportunities of intracellular replication to those able to survive in intra-protist environments. Hence, protists could serve as the training grounds and environmental reservoirs for *Rickettsiales* from which some *Rickettsiales* develop invasion mechanisms and find their niches in the animal world to become pathogenic species, as presumably the case for *Legionella* (Lamoth and Greub, 2010; Molmeret et al., 2005). It is therefore important to further investigate the diversity of *Rickettsiales* hosted by protists and how emerging pathogens might arise from there. In addition, comparative genomics studies between phylogenetically closely related *Rickettsiales* with different hosts may help identify virulence genes and provide clues to targets for vaccine development.

#### S3.4. Selection of the clock model

It is important to note that the selection of the clock model, among others, had a substantial impact on the posterior ages for both the mito- and nuclear-encoded datasets. The estimated divergence times of the *Rickettsiales* and mitochondrial lineages were consistent between using the AR and IR model implemented in MCMCTree. The IR model inferred a long branch leading to the divergence of non-*Rickettsiales* alphaproteobacteria (Fig. S4E). Consequently, the estimated posterior ages of all non-*Rickettsiales* lineages were considerably smaller if inferred with the IR model.

The analysis with mcmc3r, however, showed that the AR model outperformed the IR model, as indicated by their higher marginal likelihoods (Fig. S10A). Thus, the AR model likely better explains the rate variation across lineages in our data, and was therefore used as the preferred model. This idea is further supported by the following lines of evidence. First, molecular clock dating infers not only divergence times but also the substitution rate of each lineage (Dos Reis et al., 2016). It is well known that *Rickettsiales* and mitochondrial lineages display higher evolutionary rates than other alphaproteobacteria (Fitzpatrick et al., 2006; Martijn et al., 2018; Rodríguez-Ezpeleta and Embley, 2012; Wang and Wu, 2015). The rate differences were well estimated by the AR model but not the IR model, since under the IR model the substitution rate differences between *Rickettsiales* and non-*Rickettsiales* alphaproteobacteria, and between mitochondria and non-*Rickettsiales* alphaproteobacteria were very small (Figs. S10B, S10C). Second, the result obtained using PhyloBayes was more consistent with that obtained with the AR model than the IR model using MCMCTree (Figs. S3A, S3B). Further research is necessary to better appreciate the merits of each model in modelling the evolutionary processes on different time scales and of different species (Dos Reis et al., 2018; Ho and Duchêne, 2014).

Shu DG, Luo HL, Conway Morris S, Zhang XL, Hu SX, Chen L, Han J, Zhu M, Li Y, Chen

LZ. 1999. Lower Cambrian vertebrates from south China. *Nature* **402**:42–46. doi:10.1038/46965

Smith DR, Hua J, Lee RW, Keeling PJ. 2012. Relative rates of evolution among the three genetic compartments of the red alga Porphyra differ from those of green plants and do not correlate with genome architecture. *Mol Phylogenet Evol* **65**:339–344. doi:https://doi.org/10.1016/j.ympev.2012.06.017

Stöver BC, Müller KF. 2010. TreeGraph 2: Combining and visualizing evidence from different phylogenetic analyses. *BMC Bioinformatics* **11**:7. doi:10.1186/1471-2105-11-7

Strauss H. 1993. The sulfur isotopic record of Precambrian sulfates: new data and a critical evaluation of the existing record. *Precambrian Res* **63**:225–246. doi:10.1016/0301-9268(93)90035-Z

Szokoli F, Castelli M, Sabaneyeva E, Schrallhammer M, Krenek S, Doak TG, Berendonk TU, Petroni G. 2016. Disentangling the taxonomy of Rickettsiales and description of two novel symbionts ("Candidatus Bealeia paramacronuclearis" and "Candidatus Fokinia cryptica") sharing the cytoplasm of the ciliate protist Paramecium biaurelia. *Appl Environ Microbiol* **82**:7236–7247. doi:10.1128/AEM.02284-16

Tavaré S. 1986. Some probabilistic and statistical problems in the analysis of DNA sequences. *Am Math Soc Lect Math Life Sci*. doi:citeulike-article-id:4801403

Thomas S, Alexander W, Gilligan J, Rikihisa Y. 2016. The importance of Rickettsiales infectionsRickettsiales: Biology, Molecular Biology, Epidemiology, and Vaccine Development. pp. 3–21. doi:10.1007/978-3-319-46859-4_1

Thrash JC, Boyd A, Huggett MJ, Grote J, Carini P, Yoder RJ, Robbertse B, Spatafora JW, Rappé MS, Giovannoni SJ. 2011. Phylogenomic evidence for a common ancestor of mitochondria and the SAR11 clade. *Sci Rep* **1**:13. doi:10.1038/srep00013

Tomassone L, Portillo A, Nováková M, De Sousa R, Oteo JA. 2018. Neglected aspects of tick-borne rickettsioses. *Parasites and Vectors*. doi:10.1186/s13071-018-2856-y

Tomitani A, Knoll AH, Cavanaugh CM, Ohno T. 2006. The evolutionary diversification of cyanobacteria: Molecular-phylogenetic and paleontological perspectives. *Proc Natl Acad Sci U S A* **103**:5442–5447. doi:10.1073/pnas.0600999103

Turner EC, Kamber BS. 2012. Arctic Bay Formation, Borden Basin, Nunavut (Canada): Basin evolution, black shale, and dissolved metal systematics in the Mesoproterozoic ocean. *Precambrian Res* **208**–**211**:1–18. doi:10.1016/j.precamres.2012.03.006

Van Bel M, Diels T, Vancaester E, Kreft L, Botzki A, Van De Peer Y, Coppens F, Vandepoele K. 2018. PLAZA 4.0: An integrative resource for functional, evolutionary and comparative plant genomics. *Nucleic Acids Res* **46**:D1190–D1196. doi:10.1093/nar/gkx1002

Van Kranendonk MJ, Altermann W, Beard BL, Hoffman PF, Johnson CM, Kasting JF, Melezhik VA, Nutman AP, Papineau D, Pirajno F. 2012. A Chronostratigraphic Division of the Precambrian: Possibilities and ChallengesThe Geologic Time Scale 2012. pp. 299–392. doi:10.1016/B978-0-444-59425-9.00016-0

Van Kranendonk MJ, Philippot P, Lepot K, Bodorkos S, Pirajno F. 2008. Geological setting of Earth's oldest fossils in the ca. 3.5 Ga Dresser Formation, Pilbara Craton, Western Australia. *Precambrian Res* **167**:93–124. doi:10.1016/j.precamres.2008.07.003

Vannini C, Petroni G, Verni F, Rosati G. 2005. A bacterium belonging to the Rickettsiaceae family inhabits the cytoplasm of the marine ciliate Diophrys appendiculata (Ciliophora, Hypotrichia). *Microb Ecol* **49**:434–442. doi:10.1007/s00248-004-0055-1

Walker DH. 1996. Chapter 38 Rickettsiae In: Baron S, editor. Medical Microbiology. The University of Texas Medical Branch at Galveston.

Wang D, Yang S, Tang F, Zhu H. 2012. Symbiosis specificity in the legume - rhizobial mutualism. *Cell Microbiol.* doi:10.1111/j.1462-5822.2011.01736.x

Wang S, Meade A, Lam H-M, Luo H. 2020. Evolutionary Timeline and Genomic Plasticity Underlying the Lifestyle Diversity in Rhizobiales. *mSystems* **5**:e00438-20. doi:10.1128/mSystems.00438-20

Wang Z, Wu M. 2015. An integrated phylogenomic approach toward pinpointing the origin of mitochondria. *Sci Rep* **5**:7949. doi:10.1038/srep07949

Weinert LA, Werren JH, Aebi A, Stone GN, Jiggins FM. 2009. Evolution and diversity of Rickettsia bacteria. *BMC Biol* **7**:6. doi:10.1186/1741-7007-7-6

Werren JH, Baldo L, Clark ME. 2008. Wolbachia: Master manipulators of invertebrate biology. *Nat Rev Microbiol.* doi:10.1038/nrmicro1969

Xiao S, Knoll AH, Yuan X, Pueschel CM. 2004. Phosphatized multicellular algae in the Neoproterozoic Doushantuo Formation, China, and the early evolution of florideophyte red algae. *Am J Bot* **91**:214–227. doi:10.3732/ajb.91.2.214

Yang D, Oyaizu Y, Oyaizu H, Olsen GJ, Woese CR. 1985. Mitochondrial origins. *Proc Natl Acad Sci U S A* **82**:4443–4447. doi:10.1073/pnas.82.13.4443

Yang EC, Boo SM, Bhattacharya D, Saunders GW, Knoll AH, Fredericq S, Graf L, Yoon HS. 2016. Divergence time estimates and the evolution of major lineages in the florideophyte red algae. *Sci Rep* **6**:21361. doi:10.1038/srep21361

Yang Z, Rannala B. 2006. Bayesian estimation of species divergence times under a molecular clock using multiple fossil calibrations with soft bounds. *Mol Biol Evol* **23**:212–226. doi:10.1093/molbev/msj024

Yeoh YK, Dennis PG, Paungfoo-Lonhienne C, Weber L, Brackin R, Ragan MA, Schmidt S, Hugenholtz P. 2017. Evolutionary conservation of a core root microbiome across plant phyla along a tropical soil chronosequence. *Nat Commun* **8**:215. doi:10.1038/s41467-017-00262-8

Young JPW, Johnston AWB. 1989. The evolution of specificity in the legume-rhizobium symbiosis. *Trends Ecol Evol* **4**:341–349. doi:10.1016/0169-5347(89)90089-X

Yu G, Smith DK, Zhu H, Guan Y, Lam TTY. 2017. Ggtree: an R Package for Visualization and Annotation of Phylogenetic Trees With Their Covariates and Other Associated Data. *Methods Ecol Evol* **8**:28–36. doi:10.1111/2041-210X.12628

Yuan X, Chen Z, Xiao S, Zhou C, Hua H. 2011. An early Ediacaran assemblage of macroscopic and morphologically differentiated eukaryotes. *Nature* **470**:390–393. doi:10.1038/nature09810

Zhang Y, Golubic S. 1987. Endolithic microfossils (cyanophyta) from early Proterozoic stromatolites, Hebei, China. *Acta Micropalaeontologica Sin* **4**:1–3.
